# Expansion of OSMR expression and signaling in the human dorsal root ganglion links OSM to neuropathic pain

**DOI:** 10.1101/2025.03.26.645611

**Authors:** Juliet M Mwirigi, Ishwarya Sankaranarayanan, Diana Tavares-Ferreira, Katherin A Gabriel, Seph Palomino, Yan Li, Megan L Uhelski, Stephanie Shiers, Úrzula Franco-Enzástiga, Andi Wangzhou, Joseph B Lesnak, Samhita Bandaru, Aishni Shrivastava, Nikhil Inturi, Phillip J Albrecht, Marilyn Dockum, Anna M Cervantes, Peter Horton, Geoffrey Funk, Robert Y North, Claudio Esteves Tatsui, German Corrales, Muhammad Saad Yousuf, Michele Curatolo, Robert W Gereau, Amol Patwardhan, Gregory Dussor, Patrick M Dougherty, Frank L Rice, Theodore J Price

## Abstract

RNA sequencing studies on human dorsal root ganglion (hDRG) from patients suffering from neuropathic pain show upregulation of OSM, linking this IL-6 family cytokine to pain disorders. In mice, however, OSM signaling causes itch behaviors through a direct effect on its cognate receptor expressed uniquely by pruriceptive sensory neurons. We hypothesized that an expansion in function of OSM-OSM receptor (OSMR) in sensory disorders in humans could be explained by species differences in receptor expression and signaling. Our *in situ* hybridization and immunohistochemical findings demonstrate broad expression of OSMR in DRG nociceptors and afferent fibers innervating the superficial and deep skin of humans. In patch-clamp electrophysiology, OSM directly activates human sensory neurons engaging MAPK signaling to promote action potential firing. Using CRISPR editing we show that OSM activation of MAPK signaling is dependent on OSMR and not LIFR in hDRG. Bulk, single-nuclei, and single-cell RNA-seq of OSM-treated hDRG cultures reveal expansive similarities in the transcriptomic signature observed in pain DRGs from neuropathic patients, indicating that OSM alone can orchestrate transcriptomic signatures associated with pain. We conclude that OSM-OSMR signaling via MAPKs is a critical signaling factor for DRG plasticity that may underlie neuropathic pain in patients.

## INTRODUCTION

Neuropathic pain is an important clinical problem that is not adequately treated by any existing pharmacotherapy (*1*). While animal models give tremendous insight into pain mechanisms, they have not led to successful clinical translation to-date (*2, 3*). Clinical studies have consistently demonstrated that silencing neurons in the DRG with local anesthetic injection produces rapid and near-complete pain relief in almost all patients (*4–6*). Other studies show that DRG neurons from patients with neuropathic pain, but not those without neuropathic pain, generate spontaneous action potentials (also known as spontaneous activity or SA) originating at the soma *in vitro*, strongly implicating SA as a key mechanism driving neuropathic pain (*7, 8*). Transcriptomic studies on surgically resected DRGs from patients, and on DRGs recovered from organ donors who died with a history of neuropathic pain reveal that changes in cytokine and chemokine signaling in the DRG may play a critical role in driving the changes in sensory neuron physiology that cause neuropathic pain (*7, 9, 10*). Interestingly, some of these underlying mechanisms appear to be sex-specific (*9*), consistent with mechanistic studies in animal models (*11, 12*). A key gap in knowledge is whether these cytokines or chemokines are sufficient to produce transcriptomic, signaling or electrophysiological changes that are seen in the DRGs of individuals with neuropathic pain.

OSM is an IL-6 family cytokine that is involved in many inflammatory diseases (*13*) and has also been linked to disorders affecting the sensory nervous system such as atopic dermatitis, and prurigo nodularis (*14–17*). OSM is upregulated in itchy skin lesions (*15–19*) and mutations in the human *OSMR* gene, which encodes the OSM receptor (OSMR), cause familial primary localized cutaneous amyloidosis (FPLCA), a disorder associated with chronic itch and amyloid deposits in the dermis (*14*). These mutations cause a loss of functional signaling at OSMR leading to keratinocyte dysfunction promoting nodule formation in skin (*14*). The OSM receptor (OSMR) forms a complex with gp130 to induce signaling by OSM and can complex with IL31RA to transduce IL31 signaling (*20*). Vixarelimab is a humanized monoclonal antibody that blocks functional signaling for both IL31 and OSM by binding OSMR. Vixarelimab treatment led to relief of itch and clearing of skin lesions in a phase 2 trial of patients suffering from prurigo nodularis (*16*). While there is a clear role for OSM/OSMR in skin diseases that cause itch in humans, OSM upregulated is also found in painful tissues in diseases like rheumatoid arthritis (*21*) and OSM is strongly upregulated in the DRG of males suffering from neuropathic pain undergoing thoracic vertebrectomy surgery (*7, 9*), suggesting a broader role of this cytokine in chronic pain disorders. The primary goal of the work described here was to thoroughly characterize OSM-OSMR signaling in the human dorsal root ganglion (hDRG) to gain insight into the potential for targeting this signaling pathway for the treatment of chronic pain.

OSM shows divergence in its receptor binding and signaling mechanisms between mice, rats, and humans (*22–24*) making studying OSM signaling in a human cellular context paramount for translational relevance. In mice the cytokine binds OSMR and primarily induces JAK/STAT signaling (*22–25*). In humans OSM binds OSMR and LIFR and induces both JAK/STAT and MAPK signaling (*22–25*). Previous studies in mice show that OSM acts specifically on a distinct subset of nociceptors causing sensitization of these neurons to promote itch behaviors in mice via the JAK/STAT signaling pathway (*17*), although another study found that OSM can suppress itch behaviors in mice (*26*).

OSM is also critical for development of a subset of nociceptors in mice (*27*), and OSM treatment causes thermal hyperalgesia in mice in a manner that is dependent on nociceptor gp130 expression (*28*). In rats, where OSMR signaling more closely resembles the signaling seen at human receptors (*24*), OSM acts on a broader population of DRG nociceptors, induces MAPK signaling, and causes clear behavioral signs of nociceptive sensitization that can promote chronic pain (*29*). We hypothesized that OSM may act on multiple classes of nociceptors in humans where OSMR-mediated MAPK signaling could lead to the generation of action potentials which would cause pain in patients.

Our studies on hDRG neurons show that OSMR is expressed by several different subsets of nociceptors, with high expression in putative silent nociceptors (*30*), allowing for widespread activation of MAPK signaling upon OSM stimulation.

Electrophysiological studies show that OSM causes the generation of action potentials in hDRG neurons and transcriptomic studies indicate that OSM treatment is sufficient to induce gene expression changes that resemble those seen in the DRGs of neuropathic pain patients. We conclude that OSM/OSMR signaling is likely an important mechanism promoting chronic neuropathic pain in humans.

## METHODS

### Study Design

Our goal in this study was to first comprehensively characterize the expression landscape of OSM and OSMR in the human peripheral nervous system. The second goal was to understand transcriptional and proteomic changes resulting from OSM application to hDRG cultures and explants, and to determine whether these changes align with transcriptomic profiles hDRGs from male and female neuropathic pain patients. For each of the bulk, single-nuclei, and single-cell RNA sequencing and proteomics experiments, dissociated DRG cells were obtained from a single donor and processed in ≥3 technical replicates. The transcriptomic data from hDRGs of neuropathic pain patients used for comparative analysis were previously published (*7, 9*). Cultures were treated with OSM or vehicle for 6 hours in the transcriptomic studies, while in the proteomic studies, explant slices were treated for 3 hours. MAPK and JAK/STAT signaling assays were conducted after a 30 min incubation with the respective treatment conditions. Data analysis for all experiments except for the RNA sequencing and proteomic studies was conducted in a blinded manner with respect to experimental conditions or sex. Sample sizes for the characterization and functional assays follow what has been generally used in the field (*7, 31, 32*), with the limited availability of hDRGs also influencing our decisions. No datasets were excluded from the study.

### Study Approval

The Institutional Review Boards of UT Dallas (UTD), University of Florida, and MD Anderson Cancer Center (MDACC) reviewed and granted approval for protocols. All experiments conformed to relevant guidelines and regulations. Dorsal root ganglion tissues were collected from the donors after neurological determination of death within 2-3 hours of cross-clamp. Skin biopsies were obtained from the Department of Neurology at the University of Florida. Patients undergoing thoracic vertebrectomy at MDACC for malignant tumors were enrolled in the study, with each patient providing informed consent during enrollment. DRGs were extracted during the surgical operation and brought to the lab for further processing.

### RNAscope in situ hybridization

DRGs used for RNAScope were frozen on dry ice at the time of extraction and stored in a -80°C freezer. The human DRGs were gradually embedded in OCT in a cryomold by adding small volumes of OCT over dry ice to avoid thawing and cryosectioned at 20µm onto SuperFrost Plus charged slides. Sections were only briefly thawed to adhere to the slide but were immediately returned to the -20°C cryostat chamber until completion of sectioning. The slides were then immediately utilized for histology.

RNAscope *in situ* hybridization multiplex versions 1 and 2 was performed as instructed by Advanced Cell Diagnostics (ACD) and as previously described (*33*)The probes used were OSMR (Channel 1; Cat# 537121), *NPPB* (Channel 2; Cat# 448511-C2), *CHRNA3* (Channel 3; Cat# 482421-C3), *GFRA2* (Channel 3; Cat# 463011-C3), *FABP7* (Channel 2, Cat# 583891-C2), *CD68* (Channel 3; Cat# 560591-C3); *OSM* (Channel 1; Cat# 456381), *CALCA* (Channel 2; Cat# 605551-C2), and *P2RX3* (Channel 3; Cat# 406301-C3).

#### Tissue Quality Check

All tissues were checked for RNA quality by using a positive control probe cocktail (ACD) which contains probes for high, medium and low-expressing mRNAs that are present in all cells (ubiquitin C > Peptidyl-prolyl cis-trans isomerase B > DNA-directed RNA polymerase II subunit RPB1). All tissues showed signal for all 3 positive control probes. A negative control probe against the bacterial DapB gene (ACD) was used to check for non-specific/background label.

#### Image Analysis

DRG sections were imaged on an Olympus FV3000 confocal microscope. The raw image files were brightened and contrasted in Olympus CellSens software (v1.18), and then analyzed manually one cell at a time for expression of each gene target. Cell diameters were measured using the polyline tool. Total neuron counts were acquired by counting all of the probe-labeled neurons and all neurons that were clearly outlined by DAPI (satellite cell) and contained lipofuscin in the overlay image. For *OSMR*, *CALCA* and *P2RX3* RNAscope, we summed the neuronal counts for each target/subpopulation from all images from each section, and then calculated the population percentages for that section. We averaged the population percentages from 3 sections to yield the final population value for each donor. For *OSMR, OSM, CHRNA3, NPPB, GFRA2, FABP7* and *CD68* RNAscope we used stitched tile-scan images of hDRGs for a whole field-of-view analysis of RNA expression.

Large globular structures and/or signals that auto-fluoresced in all 3 channels (488, 550, and 647; appears white in the overlay images) were considered to be background lipofuscin and were not analyzed. Aside from adjusting brightness/contrast, we performed no digital image processing to subtract background. We attempted to optimize automated imaging analysis tools for our purposes, but these tools were designed to work with fresh, low background rodent tissues, not human samples taken from older organ donors. As such, we chose to implement a manual approach in our imaging analysis in which we used our own judgement of the negative/positive controls and target images to assess mRNA label. Images were analyzed in a blinded fashion.

### Image segmentation using Cellpose

We used a deep learning-based segmentation tool called Cellpose to automate the process of RNAscope image analysis in non-neuronal cells (*34*). Single-cell automated multiplex pipeline for RNA (SCAMPR) was used to compute the spatial gene expression in non-neuronal cells (*35*). First, images were segmented with Cellpose v2 using DAPI staining with minimal adjustments in brightness and contrast. The resulting ROI outlines were converted into FIJI ImageJ compatible ROIs for downstream processing with SCAMPR. The semi-automated signal quantification workflow was used to subtract the background and calculate image-specific threshold values. The SCAMPR FIJI Image J Macro was used to generate a Gene by Cell matrix in which the gene expression level is represented as the number of pixels expressing the gene of interest divided by the area of the ROI in pixels. The overall mean expression values for each donor image were plotted for sex difference comparisons.

### Spatial ATAC-seq hDRG data

To examine the chromatin accessibility of OSM-related genes, we used previously published hDRG spatial ATAC-seq data from Franco-Enzástiga et al., 2024 (*36*). In brief, spatial ATAC-seq was performed on 10 um hDRG tissue sections (N=3 female, N=5 male, non-pain organ donors) in AtlasXomics facilities (service code: AXO-0310). 150X150 paired-end sequencing was performed using NextSeq 2000. The sequencing depth was up to 300 million reads per sample. Processing of spatial ATAC-seq consisted of alignment of sequenced reads to the human GENCODE reference genome (v38) using Chromap. Fragment files containing coordinates of sequenced DNA molecules mapping to barcodes were generated for downstream analysis. Pixels on tissue samples were designated using AtlasXbrowser using microscopy images to create Seurat-compatible metadata files. The ArchR and Seurat (*37, 38*) packages were used to analyze data.

### Computing Euclidian distances between barcodes in 10x Visium data

Euclidean distances were calculated using SciPy (scipy.spatial.distance.cdist) in Python (version 3.8) to measure pairwise distances between neuronal barcodes and genes of interest. We then calculated the minimum, mean, and maximum relative distances of a gene of interest compared to neuronal barcodes and plotted minimum relative distance for visualization.

### Immunofluorescence (IF) Analysis of Cutaneous Innervation

Multimolecular IF analyses of normal human skin biopsies were used to detect the likely cutaneous terminations of the set of DRG neurons that have uniquely high levels of *OSMR* mRNA. These neurons are among several sets of DRG neurons that have a high level of *CALCA* mRNA and CGRP production. Based on several other studies including IF analyses of human skin biopsies, OSMR-positive cutaneous innervation was expected to be among numerous CGRP-positive “peptidergic” C-fibers that have previously been shown to terminate among several distinctly different skin structures including the epidermis, dermal papillae, papillary dermis, capillaries, sweat glands, arterioles, and hair follicles (*39–41*). This innervation has been physiologically and molecularly implicated as pruriceptive nociceptors.

Most of the IHC analyses were on 3mm punch biopsies donated by normal two male and one female siblings in their 30’s who had complete genomic profiles. Each provided pairs of bilateral mirror biopsies of hairy skin from the distal leg above the lateral malleolus and lateral mid-thigh and glabrous skin from the ulnar palmar compartment of the hand (Table S1).

Following prior published protocols (*39–44*), the biopsies were fixed for 4 hours in freshly made ice-cold 4%PFA in 0.1M phosphate buffered saline at pH7.4 (PBS), then rinsed and stored in cold PBS and sent by overnight courier to Integrated Tissue Dynamics, LLC (INTiDYN) for IHC processing and qualitative analysis following previously published INTiDYN ChemoMorphometric Analysis (ITD-CMA) SOP.

The biopsies were cryoprotected by overnight immersion in 30% sucrose in PBS, imbedded and snap frozen Optimum Cutting Temperature (OCT) gel and sectioned by cryostat at 14µm thicknesses in a plane perpendicular to the skin surface. Sections were mounted with consecutive sections rotated across 20 slides to allow for this and future multiple immunolabel combinations.

The IF analyses were supplemented with slides of unlabeled alternating sections that had been archived in the INTiDYN biobank from prior published ITD-CMA studies of identically prepared biopsies from normal control subjects that were taken from hairy skin from thoracic back dermatomes, dorsal hand, and distal leg, and glabrous skin from ulnar side palmar hand, and lateral margin of the foot (*39, 42–44*).

Following previously published protocols, the indirect immunolabeling method was used on one slide of sections from each biopsy with a rabbit monoclonal OSMR 1° antibody (Abcam # ab232684) and donkey anti-rabbit Cy3 conjugated polyclonal 2° antibody (Jackson Immuno Research #711-165-152) in combination with a mouse monoclonal PGP9.5 (UCLH1) 1° antibody (Abcam #8189), and donkey anti-mouse Alexa488 conjugated 2° antibody (ThermoFisher #A-21202). In this combination, the established pan-neuronal biomarker PGP9.5 provided a basis for assessing whether and where there was any innervation that co-labeled for OSMR as well as assessing whether OMSR was expressed on other non-innervation components of the skin. A second slide of alternating sections from each biopsy assessed OMSR labeling in combination with a polyclonal sheep CGRP 1° antibody (Abcam #195382) and donkey anti-sheep Alexa488 conjugated 2° in order to assess co-expression of OSMR and CGRP as predicted from the DRG transcriptome. Given that the primary antibodies were raised in different species, the incubations were run as a simultaneous cocktail. These and all other IF combinations below included a DAPI for IF labeling of cell nuclei.

The IF ITD-CMA also included labeling for Somatostatin (SST) because the DRG transcriptome detected uniquely high levels of SST mRNA among a distinct second set of OSMR-negative, CGRP high-expressing neurons that are implicated as peptidergic C-fiber silent nociceptors. Therefore, IF combinations were assessed on other slides of alternating sections from each biopsy utilizing combinations of a rabbit polyclonal SST 1° antibody (ThermoFisher # 17512-1-AP) with the PGP9.5 and CGRP antibodies as described above. Of particular interest was a subset of OSMR DRG neurons that also had high levels of SST mRNA and whether this supplied correspondingly unique OSMR-pos/SST-pos cutaneous innervation.

First, the above label combinations confirmed that OSMR and SST IF was detectable among subsets of cutaneous innervation, and that both completely co-labeled for CGRP. Second, both had subsets of innervation and/or other skin locations that they uniquely labeled. However, there were other sites where there was similar labeling indicative of potential co-labeling. Therefore, given that the two best OSMR and SST antibodies we identified were both rabbit polyclonals, a method documented previously used which is a two step double label procedure in which one of the 1° antibodies was incubated first followed by the donkey anti-rabbit Cy3 conjugate, and then a second incubation used the other 1° antibody followed by the donkey anti-rabbit Alexa488 conjugate. While this had the possibility that the first 1° may be double labeled with both fluorophores, anything that is uniquely labeled with the second 1° will only be labeled with Alexa488 (*45*). Therefore, another slide is incubated reversing the order of the 1° antibodies to see if that may also have double labeling and additional unique labeling. Interpreting single versus double molecular properties, such as OSMR and SST herein, is a combination of whether, where, and to what extent unique and double labeling and unique labeling occurs.

Digital images were collected using a 20X objective on an Olympus BX51-WI microscope equipped with conventional fluorescence filters (Cy3:528–553 nm excitation, 590–650 nm emission; Cy2/Alexa 488: 460–500 nm excitation, 510–560 nm emission), a Hamamatsu ER, DVC high-speed camera, linear focus encoder, and a 3-axis motorized stage system interfaced with Neurolucida software (MBF Bioscience, Essex, VT, USA). Identical standardized camera sensitivity and exposure times were used for all sections across all biopsies. Images were compiled using Photoshop routines with minimal image enhancement to show the immunolabeling as it appears under the microscope.

### Human dorsal root ganglion (hDRG) extraction and dissociation

#### University of Texas at Dallas – Price lab dissociation protocol

hDRGs were surgically extracted from the donor and immediately placed in chilled and bubbled aCSF at the local transplant site. The composition of the final working aCSF solution consisted of: 93mM N-Methyl-d-glucamine (NMDG; Sigma-Aldrich, Cat# M2004), HCl (12 N; Fisher, Cat# FLA144S500), KCl (Sigma-Aldrich, Cat# P5405), NaH_2_PO4 (Sigma-Aldrich, Cat# S5011), NaHCO_3_ (Sigma-Aldrich, Cat# S5761), HEPES (Sigma-Aldrich, Cat# H4034), d-(+)-Glucose (Sigma-Aldrich, Cat# G8270), l-Ascorbic acid (Sigma-Aldrich, Cat# A5960), Thiourea (Sigma-Aldrich, Cat# T8656), Na^+^ pyruvate (Sigma-Aldrich, Cat# P2256) MgSO_4_ (2 M; Fisher, Cat# BP213-1), CaCl_2_ dihydrate (Sigma-Aldrich, Cat# C7902), N-acetylcysteine (Sigma-Aldrich, Cat# A7250).

After being transported to the lab, the DRGs were transferred to a sterile petri dish on ice. They were further trimmed with a No. 5 forceps (Fine Science Tools, Cat#11252-00) and Bonn scissors (Fine Science Tools, Cat# 14184-09) to remove connective tissue, fat and spinal and peripheral nerve processes. The dural coats (perineum and epineurium) was also carefully pulled away, leaving only the ganglia bodies. Using Bonn scissors, the resulting DRG were further divided into approximately 2 mm thick sections. The tissue fragments were transferred to 5mL of prewarmed enzyme mix consisting of 2mg/mL of Stemxyme 1, Collagenase/Neutral Protease Dispase ((Worthington Biochemical, Cat# LS004107) and 0.1mg/mL Deoxyribonuclease I, DNAse I (Worthington Biochemical, Cat# LSOO2139) in sterile filtered HBSS (ThermoFisher Scientific, Cat# 14170161). The conical tube with enzyme mixes and tissue chunklets were placed in a shaking water bath.

For rapid turnaround cultures, the DRGS were gently triturated through a sterile fire-polished glass Pasteur pipette every 25 minutes until the tissue chunks could pass through the pipette with minimal resistance. For slow overnight dissociations, Stemxyme 1 concentrations were adjusted to 1 mg/mL and similarly to the rapid dissociation protocol, 0.1mg/mL DNAse 1 was added to sterile-filtered HBSS. This was found to be optimal for dissociating a medium to large-sized lumbar DRG. We observed that the slow enzymatic digestion method provided experimenters with greater leeway in processing tissue collected later in the day and, furthermore, led to a higher neuronal yield. After this slow enzymatic digestion step, the tissue was gently triturated with a fire-polished glass pipette to loosen any connective tissue. The dissociated DRG was passed through a 100 µm cell strainer (Corning, Cat# 431752) to remove debris and obtain a uniform cell suspension. The DRG cells were further isolated by slowly layering the cell suspension on 3ml of 10% Bovine Serum Albumin (Biopharm, Cat# 71-040) or BSA solution prepared in sterile HBSS. Multiple columns were used to improve yield if debris was found to be excessive and obstructed cell movement. The BSA gradient was centrifuged at 900 *xg* for 5min at 9 acceleration and 7 deceleration speed settings. The upper and lower phase of the gradient were aspirated. The resultant pellet was resuspended in freshly prepared and warmed BrainPhys^TM^ neuronal media (STEMCELL Technologies, Cat# 05790) supplemented with NeuroCult^TM^ SM1 (STEMCELL Technologies, Cat# 05711), N2 Supplement-A (STEMCELL Technologies, Cat# 07152), 25ng/mL recombinant human β Nerve Growth Factor or NGF (R&D Systems, Cat# 256-GF), GlutaMAX (Thermo Scientific, Cat# 35050061), and FrdU consisting of 3µg/mL 5-Fluoro-2’-deoxyuridine (Sigma-Aldrich, Cat# F0503) and 7µg/mL Uridine (Sigma-Aldrich, Cat# U3003). The density and general morphology of the neurons were assessed under a light microscope before plating on Poly-D-lysine (PDL, Sigma Aldrich, Cat# P7405) coated dishes. The cells were allowed to adhere for 3 hrs before flooding the wells with additional prewarmed media. Thereafter, the DRG cultures were maintained at 37°C and 5% CO_2_ with media changes every other day until further experimentation.

#### hDRG explant preparations

NDMG-aCSF was freshly prepared on the day of hDRG collection as described above. Upon arrival at UTD, the DRGs were further trimmed to remove excess connective tissue and sectioned on the Precisionary Compresstome VF-210-OZ to a thickness of 1mm. The acutely sliced DRG explants were allowed to recover in aCSF without NMDG for 3 hours and then transferred to serum-free BrainPhys^TM^ neuronal media supplemented with N2 and SM1 until further use.

### Immunocytochemistry and imaging

The treated cultures were immediately fixed with 10% formalin (ThermoFisher Scientific, Cat#. 23-245684) for 15 min at RT and rinsed three times with 1X PBS. 1 h block was performed at RT with 10% NGS and 0.3% Triton X-100 in 1X PBS. The coverslips were then incubated overnight at 4°C with primary antibodies chicken anti-peripherin (1:1000, EnCor Biotechnology, Cat# cpca-peri) and rabbit anti-p-eIF4E (1:1000, phospho-S209, Abcam, Cat# ab76256). Coverslips were rinsed 3 times with 1X PBS for 15 min and incubated with secondary antibodies goat anti-chicken IgY (1:2K, H+L, Alexa Fluor 647, Invitrogen, Cat# A21449) and goat anti-rabbit IgG (1:2K, H+L, Alexa Fluor 555, Thermo Fisher Scientific, Cat# A21428) for 1 h at RT. The coverslips were washed 3 times with 1X PBS and mounted with prolong gold onto uncharged glass slides and allowed to cure overnight. Slides were imaged at 20X magnification using the Olympus FV3000 RS confocal laser scanning microscope.

#### MD Anderson Cancer Center – Dougherty lab dissociation protocol

Human DRG neurons were prepared as described previously (6, 7). Excised DRG were immersed in cold (∼4°C) balanced salt solution and transported to the laboratory on ice. Each ganglion was dissected from the surrounding connective tissues and divided into sections. The sections were further divided into several ∼1-mm pieces which were placed in a petri dish containing 2 mL of a digestion solution consisting of DMEM/F12 with 0.1% trypsin (Sigma, Cat# T9201), 0.1% collagenase (Sigma, Cat# C1764), and 0.01% DNase (Sigma, Cat# D5025). The petri dish was then placed in an incubated orbital shaker at 37°C for 20 min, after which the digestion solution was collected and placed into a blocking solution (DMEM/F12 with 10% horse serum) while 2 mL of fresh digestion solution was added onto the remaining tissue pieces. This process was repeated until the tissue sections were fully digested. The combined digestion and blocking solution samples were centrifuged at 23°C and 180 *xg* for 5 min. The supernatant was then removed, and the cell pellet resuspended with 1 mL of culture media (DMEM/F12 with 10% horse serum and 1% penicillin-streptomycin). An additional 2-3 mL of culture media was added, and then the suspension filtered through a 100 µm cell strainer. The remaining suspension was centrifuged at 23°C and 180 *xg* for 5 min. The supernatant was then removed, and the cell pellet re-suspended with culture media. Cells were plated on poly-L-lysine-coated glass sheets and held in culture dishes with culture media until used (8).

### Experimental reagents

Human recombinant Oncostatin M (R&D Systems, Cat# 8475-OM or Sigma-Aldrich Cat# O9635) was preformulated and lyophilized in BSA to increase protein shelf-life. Upon receipt, it was reconstituted in 100 μg/mL of sterile PBS as per manufacturer’s guidelines and stored at -20°C for up to 3 months. Tomivosertib, also known as eFT508, was purchased from MedChem Express (Cat#HY-100022), reconstituted in DMSO at 10mM and stored at -20 °C long-term.

### Whole cell recordings of human DRG neurons

Recordings with human DRG neurons began the day following dissociation. Dissociated DRG neurons were transferred to a recording chamber perfused with extracellular solution containing 140 mM NaCl, 5 mM KCl, 2 mM CaCl_2_, 2 mM MgCl_2_, 10 mM HEPES and 11 mM glucose adjusted to pH 7.4 with NaOH. Glass-micropipettes were filled with an internal solution with 125 mM KCl, 15 mM K-gluconate, 5 mM Mg-ATP, 0.5 mM Na_2_GTP, 5 mM HEPES, 2 mM MgCl_2_, 5 mM EGTA, and 0.5 mM CaCl_2_ adjusted to pH 7.4 with KOH. The DRG neurons were held at 0 pA to record spontaneous activity for 5 min. Neurons were then held at 0 pA, and action potentials were evoked using a series of 500-ms depolarizing current injections in 10-pA steps from -50 pA. The current that induced the first action potential was defined as 1X rheobase. Only neurons with a resting membrane potential of at least -40 mV, stable baseline recordings, and evoked spikes that overshot 0 mV were used for further experiments and analysis. Series resistance (Rs) was compensated to above 70%. All recordings were made at room temperature. Whole-cell patch clamp was used to record OSM-induced effects in 6 cells from 2 patient DRGs. Human DRG neurons were monitored for resting membrane potentials for at least 5 min (baseline) until the cell had reached a steady state, and then OSM was added to the extracellular bath to achieve a concentration of 10 ng/ml. The infusion time to reach the target concentration based on the volume of the recording dish and infusion rate was approximately 2 minutes. DRG neurons were then monitored until the cessation of firing during washout of OSM.

## CRISPR

CRISPR plasmids for Cas9-mediated editing of OSMR and LIFR were constructed by GeneCopoeia (Guangzhou, China). Single synthetic guide RNA (sgRNA) clones were synthesized against all 6 variants for human OSMR (Gene ID: 9180), against all 4 variants for human LIFR (Gene ID: 3977), and a scrambled control. All sgRNA were cloned in the mammalian pCRISPR-CG12 vector with CMV-Cas9-EF1a-mCherry-IRES-neomycin in bacterial stock. Amplification of plasmid constructs was performed at UTSW.

hDRG cultures were plated at approximately 50-60% confluency on PDL-coated 6-well glass bottom plates (Fisher Scientific, Cat# NC0452316) for PCR validation or on 8-chamber culture slides (CELLTREAT, Cat# 229168) for immunohistochemistry purposes. On *DIV 3*, cells were transfected with the lipofectamine 3000 reagent (Thermo Fisher Scientific, Cat# L3000008) according to the manufacturer’s instructions and as described previously for use on hDRG cultures (*46*). Cultured cells were harvested for PCR on *DIV 5*. For signaling assays, media without transfection reagents was replaced on *DIV 5*, and on *DIV 6*, the cells were treated with OSM (10 ng/mL) for 30 minutes, followed by immunolabeling.

### Isolation and amplification of DNA

DNA lysate was collected from the cultured cells at 24h post transfection. Cells were rapidly rinsed with PBS and scraped from the plates in lysis buffer. The cells were lysed in an Accublock dry bath (Labnet International) at 65°C for 20 min and then 95°C for 10 min. The mixture was then centrifuged at 12000 rpm for 1min. The resulting DNA supernatant was collected and yield was assessed using a Nanodrop. DNA was amplified in 35 cycles.

### RNA isolation and bulk sequencing

DIV 5 hDRG cultures (approximately 1x10^5^ cells) were stimulated with OSM for 6 hours. Cells were quickly rinsed with 1X PBS twice and harvested in Buffer RLT Plus with added β-mercaptoethanol as recommended in the RNeasy Plus Micro Kit (Qiagen, Cat# 74034). Cells were lysed by vortexing for about 90s. Contaminating genomic DNA was removed following RNA isolation using a column-based approach as instructed in the kit. The yield and purity of the RNA was evaluated using a Nanodrop. RNA integrity number (RIN) was obtained using a Fragment Analyzer as a measure of RNA degradation. Ribosomal RNA was depleted from all the samples prior to generating libraries using the Illumina TruSeq RNA library prep kit V2. The resulting RNA-seq libraries were subjected to 50-cycle single-end sequencing using the Illumina Hi-Seq platform. The acquired reads were aligned to the human genome reference hg19 from the NCBI Entrez database using the bowtie2 tool. Maximum allowable alignment mismatch of 42 was used during this process. The aligned reads were calculated on StringTie as transcripts per million (TPM) for all genes while accounting for the sample library size and gene length. Differentially expressed genes between the vehicle and OSM conditions were analyzed using DESeq2 package in R.

### 10X Genomics Single-nuclei RNA-Seq sequencing

Single-nuclei RNA sequencing was performed as recommended by 10X Genomics Fixed RNA Profiling (also called the Single Cell Gene Expression Flex) protocols (*47, 48*). *DIV 4* hDRG cultures plated onto 6-well glass bottom plates were incubated with 10 ng/mL human recombinant OSM or Vehicle for 6 hours. Afterward, they were rinsed with 1x PBS and harvested. Following fixation for 16h at 4°C, we incubated the samples with the 10X Fixed RNA Feature Barcode kit for 16h. The remainder of the library preparation was done according to the manufacturer’s protocol. The samples were sequenced using NextSeq500 at the University of Texas at Dallas genome core with a pair-end reading strategy. Sequencing data were mapped to the human (GRCh38) genome and filtered using 10X Genomics Cellranger v7 (Table S2).

### PIPseq single-cell RNA sequencing

Whole-cell single-cell RNA sequencing was performed using the Particle-Integrated Polymeric emulsification sequencing, PIPseq^TM^ T2 Single Cell RNA kit (Fluent BioSciences). *DIV* 4 hDRG cultured onto 6-well glass bottom plates were treated with 10 ng/mL human recombinant OSM or Vehicle for 6 hours. Singe cells were then encapsulated in monodispersed droplets containing barcoded particle templates and lysis reagents. The PIP emulsions were processed according to the manufacturer’s instructions for mRNA isolation, cDNA generation, cDNA amplification, library preparation, library pooling, and sequencing. Sequencing of the cDNA libraries was performed on the NextSeq500 at the University of Texas at Dallas genome core (Table S2).

### Data Processing

Both 10X Genomics and PIPseq datasets were processed similarly for consistency. The count matrices were first processed using the SoupX package (*49*) to remove ambient RNA and imported into R using the Seurat package (*37, 38*). Low-quality cells/nuclei with fewer than 200 detected features or with mitochondrial content exceeding 10% were excluded from further analysis. Data were integrated using Harmony (*50*). Normalization was carried out using the SCTransform method in Seurat, which also regressed out the effects of mitochondrial gene expression. Clusters were initially annotated based on the expression of established marker genes (*51*). Low-quality clusters, including doublets, were excluded from further analysis before re-assigning final cluster identities. Differential gene expression analysis between OSM and vehicle conditions was assessed using *muscat* (*52*).

### Somalogic proteomics

Tissue lysates were prepared from acutely sliced hDRG explants treated with either 10ng/mL human recombinant OSM or vehicle for 3 hrs. Protein extraction followed our previously described protocol (*53*). The explants were placed in T-PER Tissue Protein Extraction Reagent (Thermo Scientific, Cat# 78510) with additional 1X Halt Protease Inhibitor Cocktail (Thermo Scientific, Cat# 87786) and homogenized using Precellys Soft Tissue Homogenizing beads (Bertin Corp, Cat # P000933-LYSK0-A.0). Samples were centrifuged at 14,000 *× g* for 15 minutes at 4°C. The resulting supernatant was quantified using the Micro BCA™ Protein Assay Kit (Thermo Scientific, Cat# 23235) and normalized accordingly. Proteins were profiled using the SOMAScan platform. 7000 analytes were measured on the SOMAScan assay. Quality controls were performed by SomaLogic to correct for technical variabilities within and between runs for each sample.

### Data Analysis and Statistics

Graphs and statistical analyses were generated using GraphPad Prism version 10 (GraphPad Software, Inc. San Diego, CA USA). A relative frequency distribution histogram with a Gaussian distribution curve was generated using the diameters of all *OSMR*-positive neurons detected in all experiments. Pie-chart represents the average of all of donors. Data are expressed as the mean ± SEM. Statistical analyses were performed using one-way analysis of variance (ANOVA) with Bonferroni’s multiple comparisons test, or a Wilcoxon test. A *P* value of less than 0.05 was considered statistically significant. For sequencing and proteomics results, differentially upregulated and downregulated genes and proteins were identified using R version 4.3.1 – with the muscat version 1.14.0 package for sequencing and limma version 3.56.2 package for proteomics. They were defined by a Log_2_FC ≥ 0.585 or ≤ -0.585, and an adjusted p-value < 0.05.

### Data and Code availability

Processed data will be publicly accessible at <u>sensoryomics.com</u> and through the SPARC portal. The code will be available on GitHub at https://github.com/orgs/utdal/teams/utdpaincenter.

## RESULTS

*OSMR expression in nociceptors and OSM expression in macrophages in the hDRG Osmr* expression is restricted to a distinct subset of nociceptors involved in itch behaviors in mice (*17, 26*). We sought to assess potential species differences in gene expression in humans using DRG tissues derived from organ donors. We first examined previously published datasets from hDRG (*51, 54*) and observed an expansion of OSMR expression, including at least 3 subtypes of nociceptors in humans, pruritogen receptor enriched nociceptors, putative silent nociceptors, and putative C-low threshold mechano-receptors (LTMRs) (Fig. 1A). We then used markers for these subpopulations and conducted multi-channel RNAscope *in situ* hybridization to confirm *OSMR* expression in organ donor DRGs. We observed *OSMR* expression in ∼30% of hDRG neurons, and the co-expression with gene markers of populations observed in spatial sequencing experiments was consistent (Fig. 1B and C). *OSMR*^+^ neurons co-expressed *NPPB*, consistent with pruritogen receptor enriched nociceptors, and *CHRNA3*, consistent with putative silent nociceptors (Fig. 1B-D) (*55*). A subset of *OSMR*+ neurons also coexpressed *GFRA2*, consistent with c-LTMRs (Fig. 1E-G). OSMR+ neurons also expressed broader nociceptor markers such as *CALCA* and *P2RX3* and cell size distribution was also consistent with previous observations for hDRG nociceptors (Fig. S1A-E). We examined potential sex differences in *OSMR* mRNA expression in these experiments and did not note any differences between male and female organ donor hDRGs (Fig. 1D and G, Table S1). We also noted *OSMR* mRNA puncta outside of neuronal cell bodies in hDRG (Fig. S1E). Using a marker for satellite glia cells (satglia, *FABP7*) (*56, 57*) and a marker for immune cell populations that is enriched in macrophages (CD68) (*51*), we found that *OSMR* was also expressed in satglia and immune cells in the hDRG (Fig. 1H and I, Table S1), again with no sex differences.

**Fig. 1.**
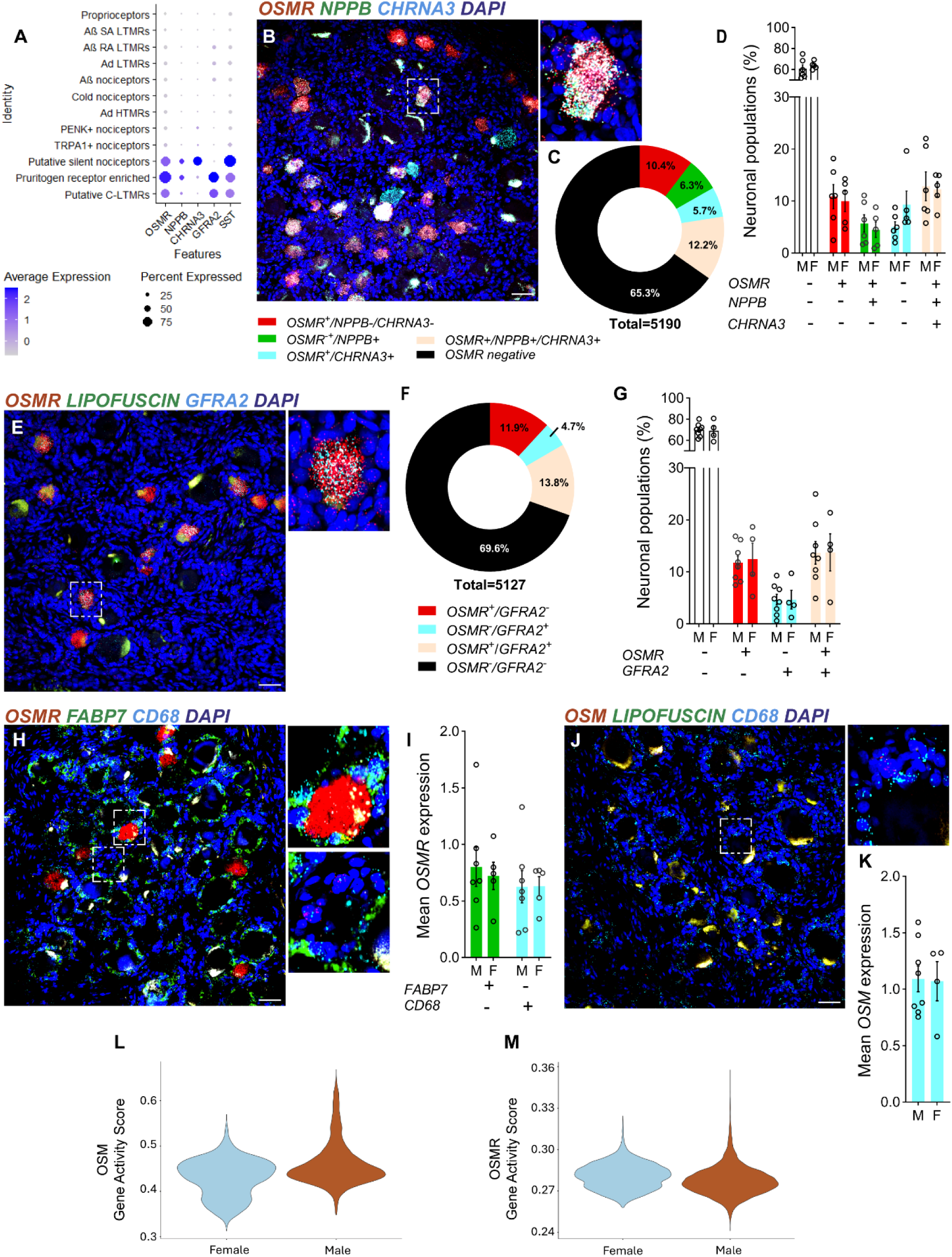
Oncostatin M (OSM) and its receptor (OSMR) are abundantly expressed in neuronal subpopulations in both male and female dorsal root ganglia from donors without pain history. A) UMAP plot of our previously published spatial transcriptomics data on human DRG (hDRG) reveals specific expression of *OSMR* in three neuronal populations: putative silent nociceptors, itch nociceptors, and C-low threshold mechanoreceptors. These neuronal subtypes expressing SST can be further distinguished by the marker genes *NPPB, CHRNA3,* and *GFRA2* (n = 4 male and 4 female donors). B) Representative RNAscope *in situ* hybridization images confirming the presence of *OSMR* mRNA puncta (red) in *NPPB*^+^ (green) and *CHRNA3*^+^ (cyan) neuronal populations. C, D) Quantification and sex-specific analysis of *OSMR* puncta in neurons with and without co-expression of *NPPB* and *CHRNA3* (n = 6 male and 5 female donors) E) Representative images showing the presence of *OSMR* (red) mRNA in *GFRA2*^+^ neurons (cyan). F, G) Quantification and sex-specific analysis of *OSMR* puncta in *GFRA2*^+^ populations (n = 8 male and 4 female donors). H, I) Detection and quantification of *OSMR* (red) in *FABP7*^+^ satellite glial cells (green) and *CD68*^+^ (cyan) macrophages (n = 7 male and 5 female donors). J, K) Representative images showing *OSM* puncta (red) are detected in a subpopulation of the *CD68*^+^ (cyan) macrophages in both sexes (n = 8 male and 4 female). L, M) Analysis of sex differences in chromatin accessibility of *OSM* and *OSMR* in hDRG using spatial ATAC-seq (n = 5 male and 3 female donors). Cell nuclei stained with DAPI (blue). 10X magnification, scale bars, 100 µm.

Previous RNA sequencing experiments on hDRG also demonstrate that OSM is expressed, but the cell type that expresses OSM in the hDRG has not been clarified. Using RNA scope *in situ* hybridization we observed that *OSM* was expressed in CD68+ cells, likely macrophages (Fig. 1J and K). To confirm the expression and lack of sex differences findings with an independent method, we also analyzed a previously published spatial ATAC-seq dataset of hDRG (*58*). We examined open chromatin around the OSM and OSMR genes finding gene accessibility for both genes, but no sex differences in open chromatin peaks for either gene (Fig. 1L and M).

Finally, we used previously published spatial hDRG RNA sequencing (*54*) to quantify the arrangement of *OSMR* and *OSM*-expressing cells in the hDRG. Calculating the Euclidian distances between barcodes expressing these genes revealed that OSM-expressing cells were arranged in close proximity to neurons expressing the receptor, suggesting a physiological relationship between these cells (Fig. S2A and B). In humans, OSM can also bind to LIFR, therefore we also evaluated *LIFR* expression in previously published spatial and single nucleus sequencing datasets from hDRG. Unlike *OSMR*, *LIFR* was ubiquitously expressed in hDRG neuronal populations and was also expressed in most other cell types (Fig. S3A-F).

### OSMR localizes to afferent terminals of the human epidermis and deeper skin layers

Having established *OSMR* mRNA expression in specific populations of hDRG neuronal cell bodies, we assessed OSMR expression in human skin biopsies to determine if protein could be detected in nerve terminals expressing markers for specific cell types. OSMR overlapped with populations that also expressed *SST*, and a subset of CGRP-expressing human nociceptors that are likely silent nociceptors (*30, 59, 60*). OSMR and SST IF were detected as subsets among all the cutaneous innervation revealed by anti-PGP9.5 staining (Fig. 2A-D; Fig. S4A, B,E, and Fig. S5A-D). Both OSMR and SST consistently co-labeled for CGRP, aligning with sequencing data showing that these populations should overlap in humans (Fig. 2E-H and Fig. S4C,D,F) (*30, 54, 59*). However, CGRP was also consistently expressed among innervation that was OSMR and SST-negative. SST IF was detected only on nerve terminals with additional very low-level expression on keratinocytes (Fig. 2A,B,E,F; Fig. S4A,C; Fig. S5A,B,E,F). OSMR IF was not only detectable among nerve terminals, but also epidermal keratinocytes, terminal glia in Meissner corpuscles, arteriole smooth muscle, sweat glands, and scattered cells in the dermis (Fig. 2C,D,E,H; Fig. S4E,F; Fig. S5C,D,G,H).

**Fig. 2.**
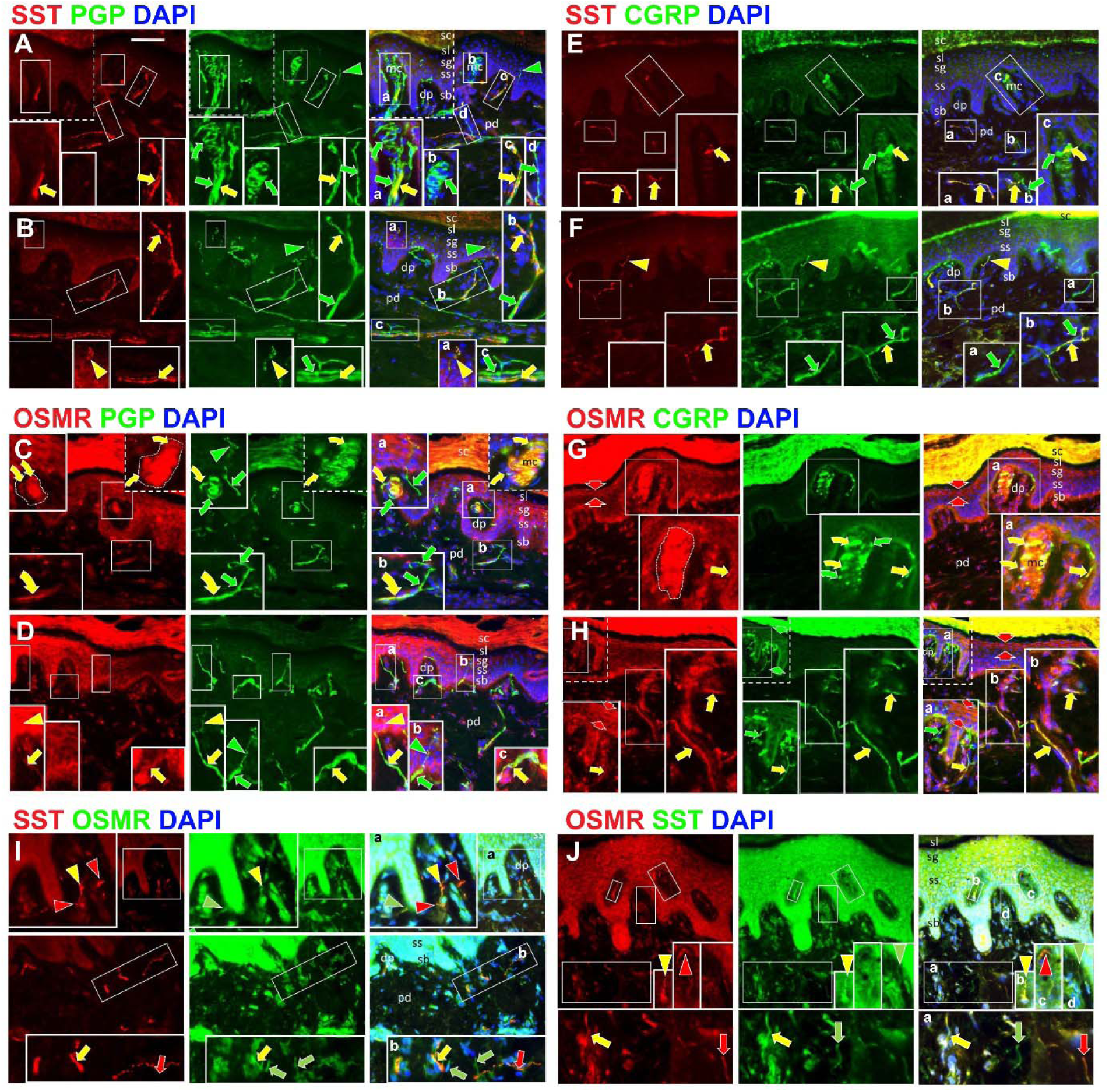
Immunofluorescence labeling of human palmar skin biopsies reveals the expression of OSMR and SST in small nerve fibers, free nerve endings (FNE), terminal glia, and keratinocytes. IF labeled antigen combinations are noted above uppercase lettered panels for separate red and green IF channels (left and center images) and for merged IF (right images) with cell nuclei co-labeled for DAPI in blue IF. Epidermal layers of keratinocytes from deep to superficial are: sb, stratum basalis; ss, stratum spinosum; sg, stratum granulosum; sl, stratum lucidum; sc, stratum corneum. Arrowheads indicate IF sensory endings in the epidermis and invaginating dermal papillae (dp). Straight arrows indicate axons and endings in the collagen-dense papillary dermis (pd) subjacent to the epidermis. Arrowheads and arrows are red or green, indicating innervation detected by IF for only one corresponding antigen, and yellow for co-labeling. Scale bar in A, 50µm. Larger solid line rectangles in each image are 2X enlargements of locations in smaller rectangles as indicated by corresponding lower-case letters in the right images of each panel. Dash-lined rectangles are inserts taken from images of other sections. **A-D**) SST and OSMR are respectively expressed as subsets among the total innervation labeled for PGP9.5 (yellow indicators) whereas other innervation is SST and OSMR negative (green indicators). **E-H**) All SST and OSMR fibers are co-expressed with CGRP (yellow indicators), but other CGRP fibers are SST or OSMR negative (green indicators). Note that SST is expressed virtually only on innervation whereas OSMR is also expressed on keratinocytes, sometimes mostly in sg (**G, H** between broad red arrows), as well as among other cells in the dermis (small white arrows). On occasion, SST can be detected in endings that definitively terminate in the epidermis (**B, inset a**) but OSMR IF on keratinocytes obscures whether OSMR is expressed on endings that penetrate the epidermis. **C,G**) OSMR is expressed on terminal glia of Meissner corpuscles (mc) as well as a subset CGRP endings that terminate on the glia (curved arrows). **I,J**) sequential double labeling as described in the methods, first with SST followed by OSMR (**I**) and then first with OSMR (**J**) reveals axons and endings and axons that only express SST, only OSMR, or SST and OSMR.

The OSMR, SST, PGP9.5, and CGRP IF combinations (Fig. 2I,J) revealed that OSMR and SST immunolabeling was entirely co-expressed among the CGRP C-fibers and their terminals within the papillary dermis and dermal papillae. However, OSMR and SST was mostly not co-expressed and likely on separate sets of CGRP fibers. The proportions of OSMR and SST positive fibers was qualitatively similar. Still other CGRP fibers were both OSMR and SST negative. Since, the keratinocytes robustly expressed OSMR, it was difficult to detect whether OSMR was expressed among the CGRP innervation that terminates in the epidermis. However, OSMR was expressed on fibers that were in contact with the basement membrane and likely to terminate in the epidermis (Fig. 2D). Whereas, keratinocytes had no confounding SST labeling, SST was detected on some intraepidermal nerve fibers (IENFs, Fig. 2B inset a) but, for the most part, was not seen on SST-positive PGP9.5 labeled fibers whose endings continued into the epidermis. Unlike in rodents and even monkeys where CGRP-IF is readily express on ∼10% of IENF supplied by CGRP-positive fibers in the subjacent dermis (*61, 62*), IENF in human skin biopsies rarely label for CGRP even though it is expressed in the upper dermal fibers up to but not beyond where they penetrate the epidermis as seen by PGP9.5-IF.

As shown in insets in Fig. 2C,E, and G, many dermal papillae in glabrous skin contain a Meissner Corpuscle (mc) which contain stratified terminals from one or more Aβ fibers that are intermingled with terminals from C-fibers. OSMR IF was expressed on the terminal glia and likely also the CGRP-positive terminals (Fig. 2C,G). SST was rarely expressed on these CGRP+ terminals (Fig. 2E).

Previous publications have documented that arterioles and arteriole-venule shunts have the highest innervation density in the skin of humans and most mammalian species (*41, 44*). Located in the relative low collagen fiber density reticular dermis deep to the papillary dermis, the cutaneous arterioles and arteriole-venule shunts are composed of three layers (Fig. S4E,F insets). The lumen is lined with the tunica intima composed of a single layer of flat endothelial cells, which is surrounded by a thick tunica media composed of several layers of circumferentially and diagonally oriented smooth muscle. The tunica media is in turn surround by an outer collagen fiber dense tunica adventitia, which is where all of a dense innervation terminates which is stratified by different types. First, noradrenergic CGRP-sympathetic terminals are concentrated near the tunica media whereas sensory endings supplied overwhelmingly by C-fibers and a few Aδ are distributed throughout the epidermis of which nearly all are CGRP+ (39, 42). Of particular interest herein, nearly all of the CGRP+ innervation to the outer half of the tunica adventitia co-labeled only for OSMR and a small proportion only for SST (Fig. S4D). However, most of the innervation to the inner half co-express both OMSR and SST (Fig. S4E,F). A small portion of CGRP fibers lacked OMSR or SST labeling (Fig. S4C, D).

Also located in the reticular dermis, sweat glands are prevalent in the reticular dermis of both human glabrous and hairy skin. Previous studies had shown that they are innervated by extensive cholinergic sympathetic fibers and some intermingled CGRP+ C fibers (*41*). OSMR was expressed only among a subset of the CGRP innervation (Fig. S4G). SST labeling was extremely rare (Fig. S5A,B,E,F). Consistent with the innervation observed at all levels of the skin, the dermal nerves located in the reticular innervation contained a mix of fibers of which OSMR and/or SST were only expressed among CGRP-positive fibers. Among these some were only OSMR-positive, some were only SST-positive, some were positive for both OSMR and SST, while others were negative for OSMR and SST.

Overall, these results support the conclusion that OSMR is expressed in multiple populations of human C fibers and is found in nerve terminals innervating multiple areas of human skin suggesting that these nerve fibers are able to detect stimuli superficially, but also at deeper structures in the skin consistent with a role in both pain and itch and broad expression in what are potentially dermal silent nociceptors (*30*).

### OSM directly activates hDRG neurons via MAPK-MNK1/2 signaling

Having established OSMR expression in hDRG neuron cell bodies and their peripheral terminals in the human skin, we turned to understanding the effect of OSM on hDRG neurons *in vitro*. We cultured hDRG neurons recovered from organ donors and after 4 days *in vitro* (DIV) stimulated with increasing concentrations of OSM (3-30 ng/mL). We used the target of MNK signaling, eIF4E phosphorylation at Serine 209, as a functional readout since OSM is known to stimulate MAPK signaling in certain cell types (*63, 64*) and eIF4E phosphorylation by MNK is a specific biochemical readout of this signaling pathway, and MNK-eIF4E signaling has been extensively linked to nociceptor sensitization in rodent and human studies (*31, 32, 50, 65–70*). We observed a concentration-dependent increase in eIF4E phosphorylation with OSM treatment for 30 min (Fig. 3A-E). This effect was completely blocked by the specific MNK inhibitor eFT508 (25 nM, Fig. 3F-J). To test OSM-induced MNK signaling in hDRG neurons with another technique we made acute slices from DRGs recovered from organ donors and stimulated these explants with 10 ng/mL OSM. We used immunofluorescence for p-eIF4E to assess signaling in specific cells in intact hDRG slices. Consistent with culture experiments, we observed increased p-eIF4E in neuronal cell bodies (Fig. S6A-C), but we also detected increased p-eIF4E around neurons consistent with expression of OSMR in satglia (Fig. S6D).

**Fig. 3.**
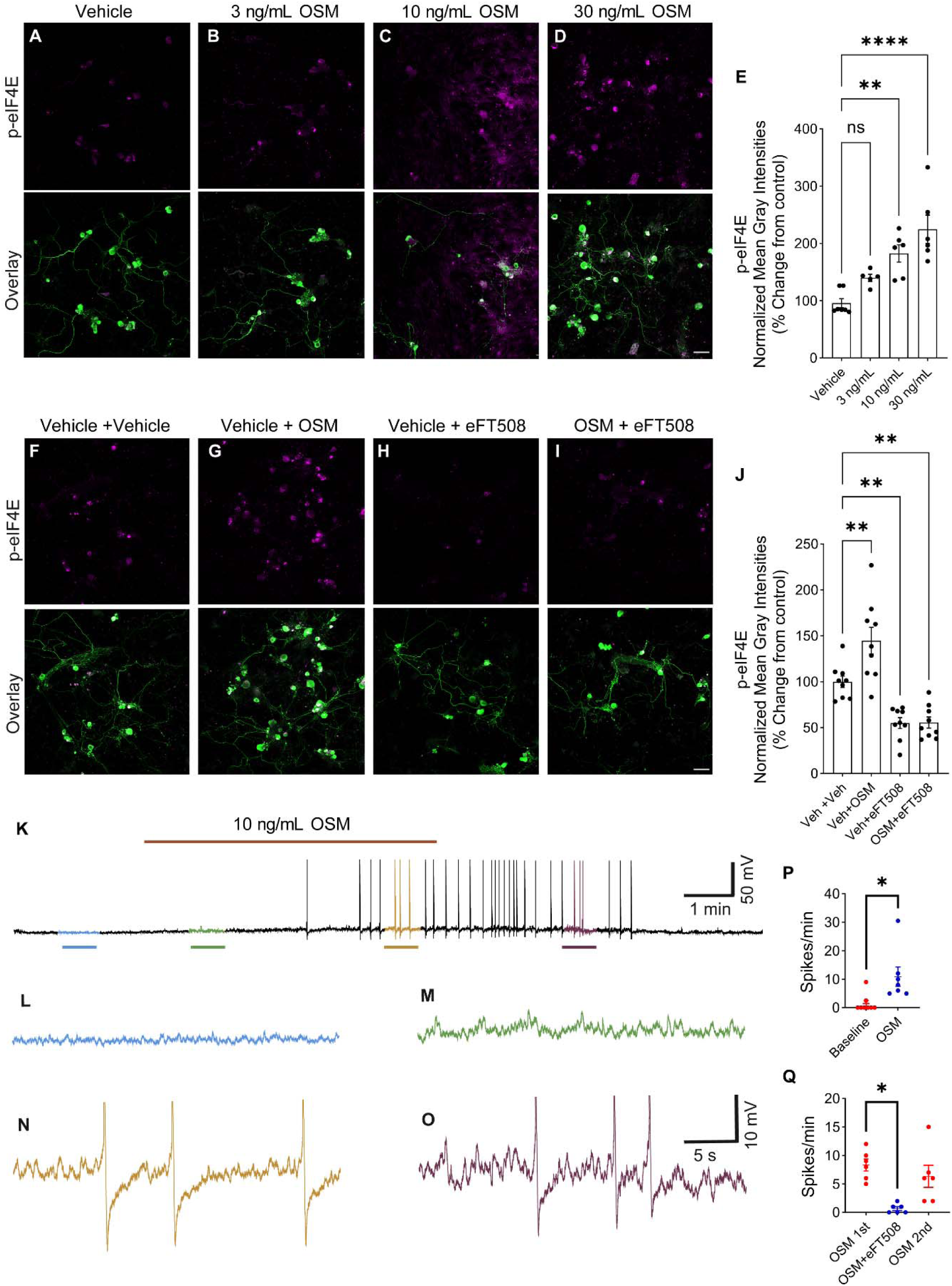
OSM activates the MNK-eIF4E signaling axis in a concentration-dependent manner and induces spontaneous activity in cultured hDRG neurons, an effect blocked by eFT508. **A-E**) 3, 10, and 30 ng/mL human recombinant OSM or Vehicle were applied to applied to DIV 4 hDRG cultures for 30 min. Representative confocal images show immunostaining with phospho eIF4E (p-eIF4E, purple) in peripherin-labeled neurons (green). 10X MagnificaScale bar, 100 µm. Normalized mean gray intensities illustrate a significant increase in p-eIF4E signal intensities at 10 and 30 ng/mL OSM treatments. N = 7, 5, 6, 6 technical replicates for Vehicle, 3 ng/mL, 10 ng/mL, 30 ng/mL OSM respectively. Cultures were prepared from lumbar DRGs from 2 donors. **F-I**) Representative confocal images showing p-eIF4E with and without OSM and eFT508 treatment for 30 min. OSM-induced increases in p-eIF4E signal intensities were reversed by MNK1/2 inhibitor eFT508 (25 nM). N = 9 technical replicates for Vehicle + Vehicle, Vehicle + 10 ng/mL OSM, Vehicle + eFT508, and OSM+ eFT508 conditions. Cultures were prepared from lumbar DRGs from 2 donors. Data represent means ± SEM. Ordinary one-way ANOVA with Bonferroni’s posttest: \*\**P* < 0.01, \*\*\**P* < 0.001. Scale bars, 100 µm. **K-L**) A representative raw trace demonstrating the onset of action potentials during OSM application. **M-N**) Representative trace of a neuron firing action potentials after an initial OSM application, which slows after a washout period. Another application of OSM triggers action potential discharge once again, but after co-treatment with eFT508, action potential firing is suppressed. After a washout period, a second treatment with OSM alone results in an action potential firing rate similar to the first OSM treatment. Data represent means ± SEM. Wilcoxon test: \**P* < 0.05.

In mice, OSM does not directly excite nociceptors but sensitizes them to itch mediators (*17*). Using patch-clamp electrophysiology on hDRG neurons recovered from organ donors we found that OSM directly excites DRG neurons causing them to fire action potentials that commence shortly after OSM (10 ng/mL) exposure (Fig 3K). OSM-induced action potential firing was preceded by instability in the resting membrane potential resembling what has been described in rodent and human nociceptors from neuropathic pain models and patients (*71–79*), respectively, and referred to as depolarizing spontaneous fluctuations (DSFs) (Fig. 3K-M). This was followed by firing of action potentials that continued even after OSM was removed from the bath (Fig. 3N-P). The effect of OSM was blocked by eFT508 treatment, indicating that MNK-eIF4E signaling drives this change in excitability (Fig. 3Q). These experiments show that OSM directly excites hDRG neurons, illustrating another species difference in the actions of this cytokine on nociceptors.

In addition to MAPK signaling, OSM also activates STAT signaling in many cell types (*64, 80–82*). Therefore, we also examined whether OSM increases STAT3 phosphorylation in hDRG neurons 4 DIV. We observed a concentration-dependent increase in p-STAT3 in hDRG neurons in response to OSM treatment (Fig. 4A-E). While we attribute the excitability changes in hDRG neurons induced by OSM to MAPK signaling (Fig. 3), changes in STAT3 activity suggest that OSM may also cause transcriptional changes in hDRG cells that can be assessed with RNA sequencing experiments.

**Fig. 4.**
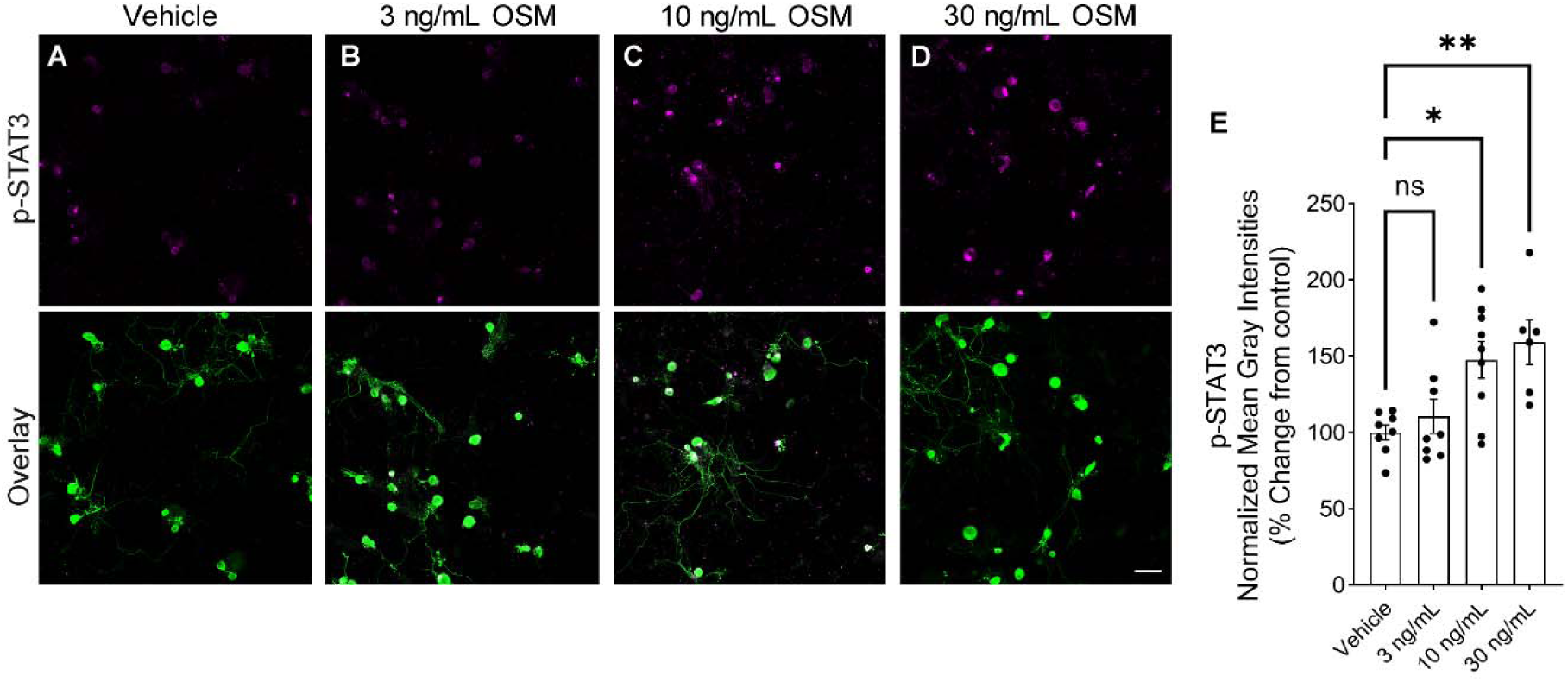
OSM activates the STAT3 signaling in a dose-dependent manner cultured hDRG neurons. **A-D**) 3, 10, and 30 ng/mL human recombinant OSM or Vehicle were applied to applied to DIV 4 hDRG cultures for 30 min. Representative confocal images show immunostaining with phospho STAT3 (p-STAT3, purple) in peripherin-labeled neurons (green). p-STAT3 signal intensities are significantly increased application of 10 and 30 ng/mL OSM. Data represent means ± SEM. Ordinary one-way ANOVA with Bonferroni’s posttest: \**P* < 0.05, \*\**P* < 0.01. 10X Magnification. Scale bar, 100 µm.

### CRISPR editing shows that OSMR and LIFR mediate different actions of OSM in hDRG

Human OSM acts via two receptors that can both complex with gp130 to induce cellular signaling, OSMR and LIFR (*24*). Specific antagonists of these receptors have not been described, making it challenging to determine which receptors contribute to the actions of OSM on hDRG neurons. To assess this, we turned to CRISPR editing of OSMR and LIFR in hDRG neurons using a recently described approach of plasmid transfection of CRISPR enzymes and guide RNAs into hDRG neurons *in vitro* (*46*).

Transfection with plasmids led to robust expression of mCherry reporter in hDRG neurons and other cell types (Fig. 5A and B, Fig. S7 A-C), as expected. Knockout of OSMR led to a complete loss of OSM-induced p-eIF4E increase but this effect was intact in the negative control, and in cells where LIFR was knocked out (Fig. 5A-C). Therefore, CRISPR knockout of these receptors demonstrates that OSMR transduces OSM-mediated increases in MNK signaling to eIF4E.

**Fig. 5.**
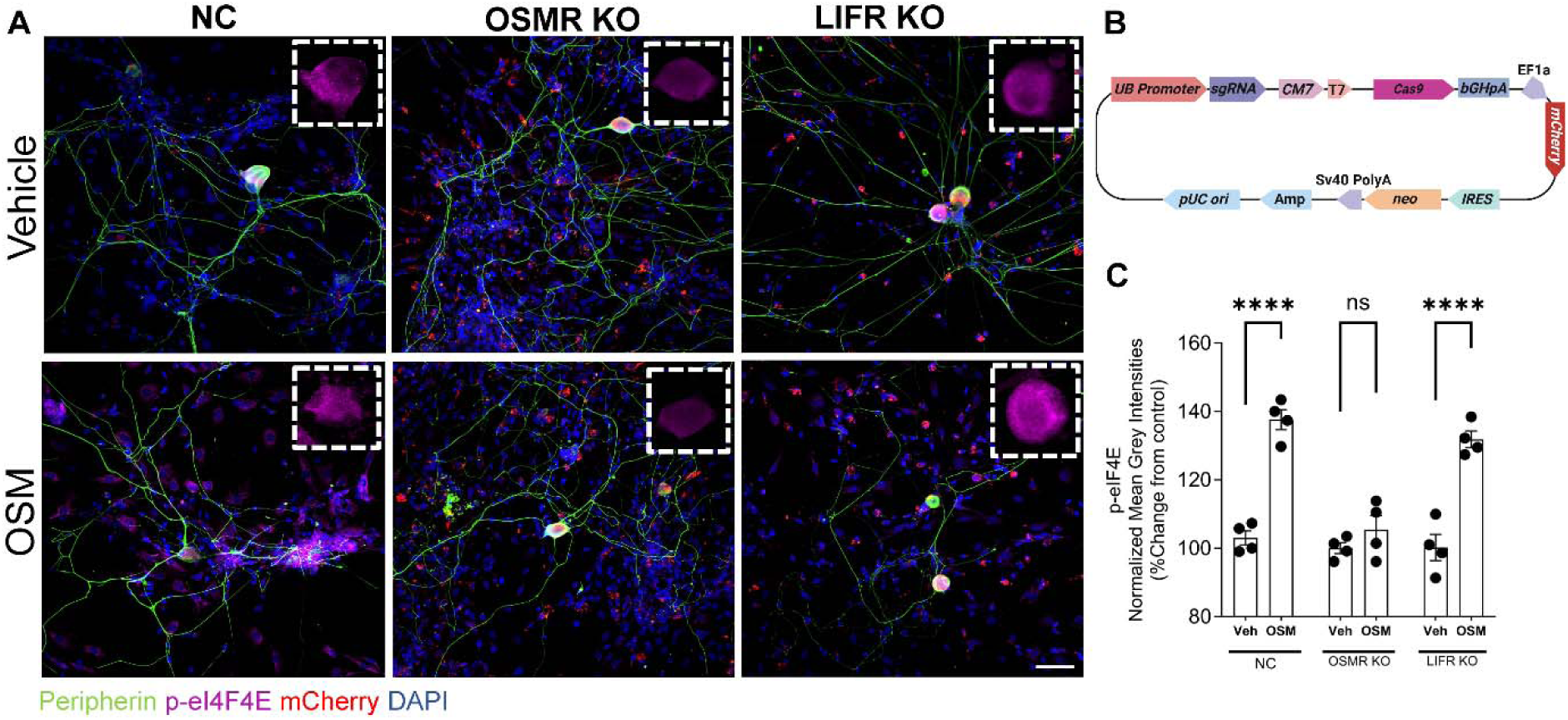
OSM-induced increases in p-eIF4E signaling are OSMR and *not* LIFR-mediated. **A**) The upper panel corresponds to the vehicle-treated while the bottom panel shows OSM-treated groups (10 ng/mL, 30min). Dashed white boxes highlight the zoomed-in neuronal cell bodies. **B**) The pCRISPR-CG12 construct used for knocking out OSMR or LIFR includes a U6 promoter for guide RNA expression, a CMV promoter for Cas9 expression, an mCherry reporter, a neomycin resistance gene for selection, and an IRES for efficient translation of the resistance gene. **C**) OSM significantly increases phosphorylation of eIF4E in the NC and LIFR KO groups but not in the OSMR KO group. Data represent means ± SEM. N = 4 replicates per cohort. Ordinary one-way ANOVA with Bonferroni’s posttest: \*\*\*\**P* < 0.0001. 10X Magnification. Scale bar, 100 µm.

### OSM causes transcriptional reprogramming of hDRG consistent with findings in neuropathic pain DRGs

Our previous work demonstrates that OSM upregulation is associated with neuropathic pain in thoracic vertebrectomy patients (*7, 9*). To understand the impact of OSM on hDRG neurons and other cell types in the ganglia, we generated mixed cultures from organ donors with no history of chronic pain and treated these cultures at DIV 4 with OSM (10 ng/mL) for 6 hrs. We reasoned that this time course would be sufficient to induce transcriptional changes in cells expressing OSMR and/or LIFR, and also to induce network changes that might be caused by factors released by cells in response to OSM. First, we isolated total RNA from the mixed hDRG cultures and did bulk RNA sequencing on 4 vehicle and 4 OSM independently treated culture wells from a single donor. The bulk RNA sequencing results revealed that OSM induces clear transcriptional responses, as evidenced by the distinct clustering of OSM-treated and vehicle-treated samples on the PCA plot (Fig. 6A). Several cytokines and chemokines were significantly upregulated, including *IL6*, *LIF*, *OSM*, *CXCL1*, *CXCL3*, *CXCL13*, *CCL2*, and *CCL7* (Fig. 6B), many of which are strongly implicated in driving JAK/STAT and MAPK signaling, sensitizing nociceptors, and contributing to the activation of a pro-inflammatory milieu (*83–85*). *SOCS3*, a potent suppressor of JAK/STAT signaling, was one of the most enriched genes, likely functioning as an intrinsic mechanism to control the excessive activation of inflammatory pathways driven by cytokine-induced responses (*86*). Additionally, *IL24* and *CCL11*, which showed increased expression and are less well-characterized in the context of pain, may play roles in modulating immune responses and neuronal-glial interactions. Key pro-nociceptive ligands and vascular remodeling factors, *VEGFA* and *ANGPTL4*, were also significantly elevated in OSM-treated samples (*87*). Additionally, neuronal-associated genes like *TAC1*, *RAMP1*, *OSMR*, *OXTR*, and *CACNB3*, alongside circadian regulators *BMAL2* and *GPR176*, potentially linking these pathways to diurnal pain fluctuations. Overall, our results demonstrate that OSM profoundly influences multiple pain-associated pathways (Fig. 6D), emphasizing its role in shaping the transcriptional landscape of pain signaling.

**Figure 6.**
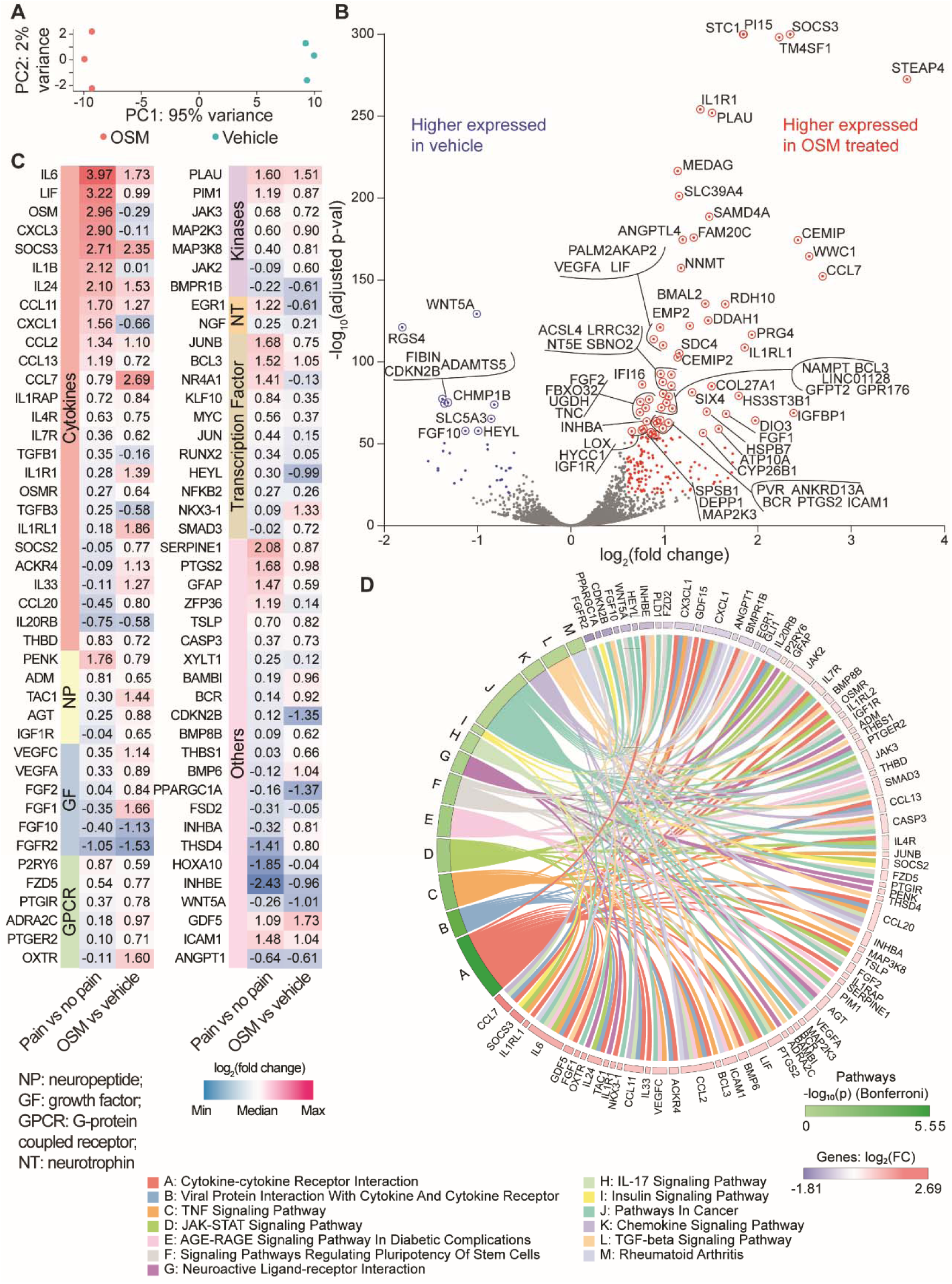
OSM drives gene expression changes in hDRG cultures, recapitulating key transcriptomic signatures found in human neuropathic pain DRGs. Bulk sequencing was performed on DIV4 hDRG cultures treated with 10 ng/mL OSM for 6 hours. **A**) Principal component analysis (PCA) plot demonstrates a separation between the OSM-treated and vehicle-treated groups, indicating distinct transcriptional profiles in response to OSM. N = 3 replicates per condition. **B**) Volcano plot of differentially expressed genes. Red dots represent upregulated genes (Log2FC ≥ 0.585) and blue dots represent downregulated genes (Log2FC ≤ -0.585) with adjusted p-value < 0.05. Labeled genes are circled. **C**) Comparison of transcriptomic data from OSM-treated cultures and neuropathic pain DRGs reveals shared gene expression profiles across GPCRs, cytokines, transcription factors, kinases, neuropeptides, and growth factors. **D**) Circos plot links shared differentially expressed genes to their enrichment in multiple signaling pathways.

We extended our analysis to directly compare OSM-treated bulk RNAseq datasets with previously published profiles of DRGs from neuropathic pain patients (*7, 9*) to identify shared and divergent gene sets that may drive nociceptor sensitization and immune response modulation. Our integrative analysis revealed substantial overlap in the upregulation of proinflammatory cytokines such as *IL6*, *LIF*, *OSM*, *SOCS3*, *IL24*, and chemokines including *CCL11*, *CXCL1*, *CCL2*, *CCL13*, *CCL7*, alongside their respective receptors *IL1RAP* and *IL4R*. We also identified increased expression of neuropeptides (*THBD*, *PENK*, *ADM*) and G protein-coupled receptors including *P2RY6*, *FZD5*, and modestly *ADRA2C*. Key kinases, including *JAK3*, *MAP2K3*, *MAP3K8*, *PLAU*, and *PIM1*, were upregulated, along with downstream transcription factors such as *JUNB* and *BCL3*, paralleling patterns observed in neuropathic pain samples (*7, 9*). These findings suggest that OSM-induced transcriptional changes recapitulate critical aspects of the transcriptional landscape seen in neuropathic pain conditions (Fig. 6C and D).

Our findings were consistent across sexes, indicating that differences in cell type proportions likely drive most sex-specific mechanisms in neuropathic pain rather than intrinsic differences in receptor biology and signaling.

To understand the impact of OSM treatment at the single-nuclei level, we conducted a second experiment using a culture from an independent donor, maintaining the same number of culture wells per treatment (4 vehicle vs. 4 OSM). As expected, neuronal nuclei were sparse; however, we successfully captured a diverse array of other cell types commonly found in native human DRG (hDRG) tissue and hDRG cultures. We identified 61,749 nuclei and annotated cell types based on their established marker genes (Fig. 7A-D, and Fig. S8A): satglia (enriched for *KCNJ10*, *GLUL*, and *FABP7*), myelinating Schwann cells/Schwann_M (expressing MPZ and MBP), and non-myelinating Schwann cells/Schwann_N (expressing SCN7A). Cell clusters also included Fibroblasts (*DCN*, *PDGFRA*, *MGP*), macrophages (*CD163*, *CSF1R*, *CCR2*, *MRC1*, *LYZ2*), B cells (*CD79A*, *CD79B*, *IGHD*, *IGHM*, *MS4A1*), T cells (*CD3E*, *CD8A*, *CCR7*, *CD4*), endothelial cells (*PECAM1*, *CLDN5*, *EGFL7*), and dendritic cells (*FCER1A*, *CST3*). We performed differential expression analysis using the muscat R package, which revealed subpopulation-specific transcriptomic changes induced by OSM. Among the identified cell types, satglia and fibroblasts exhibited the most pronounced differential changes in their transcriptomic profiles, based on a Log2FC ≥ 0.585 or ≤ -0.585 and an adjusted p-value (p_adj.loc) ≤ 0.05. Additionally, we observed the highest degree of overlap in transcriptional changes among the satglia, Schwann_M, and Schwann_M subpopulations, suggesting that these glial cell types share common signaling pathways and exhibit relatively similar cell state responses to OSM. Osteopontin (SPP1), significantly increased in satglia, regulates neurite outgrowth, mechanosensory thresholds, and knee pain via integrin-mediated ERK1/2 signaling. Neuregulin 1 (*NRG1*) overexpression in satglia suggests neuron-glia communication via ErbB receptor tyrosine kinase signaling, modulating neuronal excitability and facilitating repair processes. Ion channel and receptor genes like *TRPM3* and *P2RX7* were significantly decreased in satglia. Several genes, including *SOCS3*, *CXCL12*, *JUNB*, *PTX3*, *LMCD1*, *MAP3K8*, *IL33*, *ANGPTL4*, *GGCT*, *HAS2*, and *NAMPT*, involved in inflammatory signaling, transcriptional regulation, and cellular stress responses, were upregulated in satglia and fibroblasts, corroborating bulk RNAseq findings and highlighting satglia- and Fibroblast-specific mechanisms (Fig. 7F and G). We also observed that several members of the SERPINE family, such as *SERPINE1*, *SERPINE3*, *SERPINEB3*, and *SERPINEB4*, were notably upregulated in Fibroblasts, indicating their involvement in a profibrotic cell state characterized by enhanced extracellular matrix remodeling, and inflammatory signaling (Fig. 7G).

**Figure 7.**
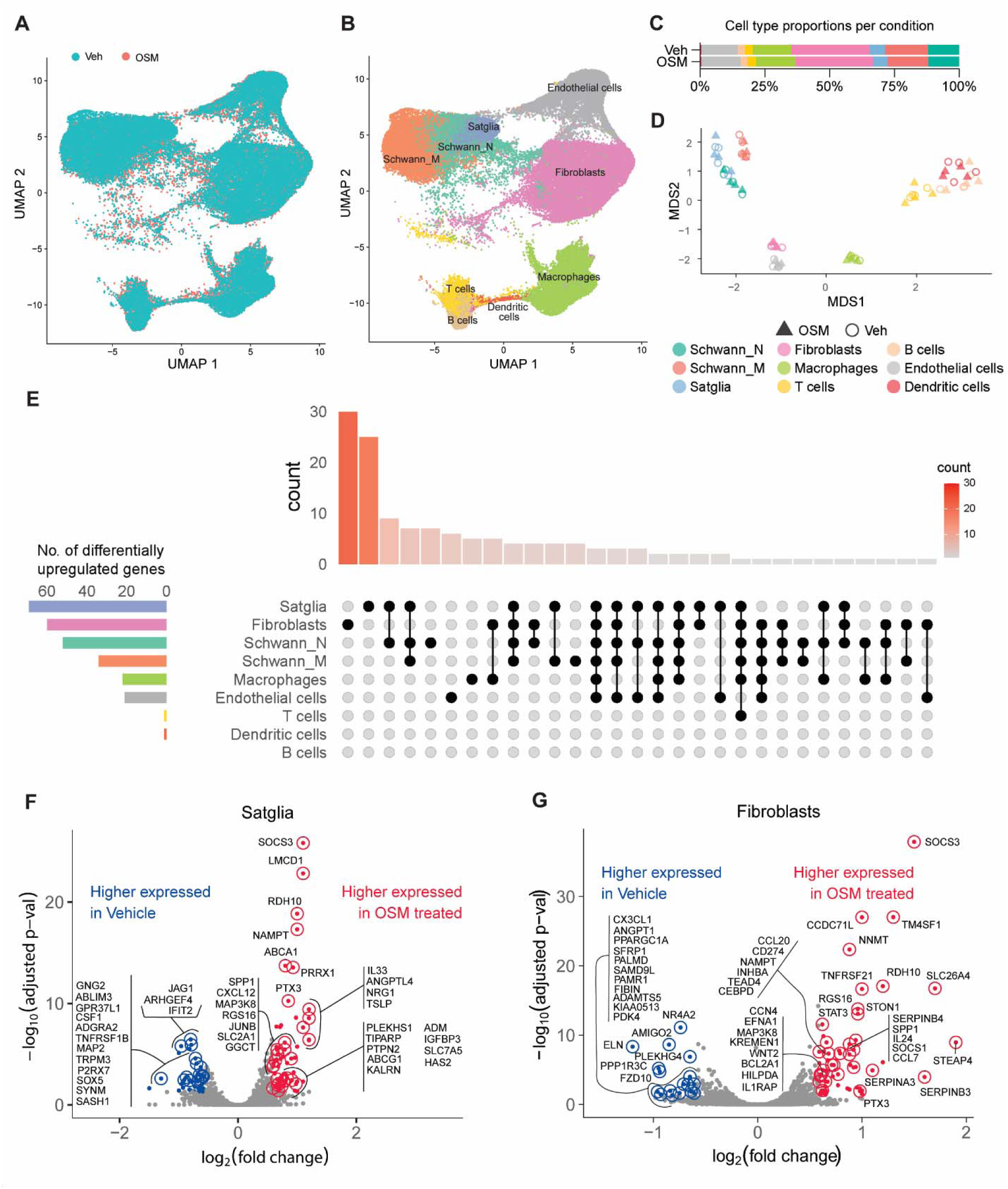
OSM elicits cell-type-specific transcriptional changes in hDRG cultures identified by sn-RNAseq. **A**) UMAP projections of OSM and Vehicle conditions, N = 4 replicates per condition. DIV4 hDRG cultures were treated with 10 ng/mL OSM for 6 hours. **B**) UMAP projection depicting the clustering of nine identified non-neuronal cell populations within hDRG cultures. Major cell types identified include satglia, fibroblasts, Schwann cells, macrophages, endothelial cells, dendritic cells, and immune cells, all captured via snRNA-seq. **C**) Proportions of each cell type in OSM-treated and vehicle-treated groups **D**) Multidimensional scaling (MDS) plot illustrating the transcriptional variation between OSM-treated and vehicle-treated samples for each cell type **E**) Upset plot displaying the number of upregulated genes across different cell types following OSM treatment. satglia and fibroblasts exhibit the most transcriptional changes, with the plot indicating both genes unique to individual cell types and those shared between multiple cell types. **F**) Volcano plot of differentially expressed genes in satglia cells, with upregulated genes (Log2FC ≥ 0.585, adjusted p-value < 0.05) shown in red, and downregulated genes (Log2FC ≤ -0.585) in blue. **G**) Volcano plot for fibroblasts, highlighting genes upregulated in OSM-treated fibroblasts (red) and downregulated in vehicle-treated fibroblasts (blue), based on the same significance thresholds.

Finally, we used an independent method, PIP-seq, to capture whole cells from hDRG cultures. Cultures were generated from an independent organ donor and treated with either vehicle or OSM for 6 hours at 4 days in vitro (DIV). While we again obtained only a small number of neurons, we successfully captured representative populations of other cell types present in the hDRG. Using the same markers, we classified 14,187 cells into satglia, Schwann_N, Schwann_M, fibroblasts, pericytes, endothelial cells, dendritic cells, and macrophages (Fig. 8A-D and Fig. S8B). As expected, the scRNA-seq approach detected more differentially expressed genes across all subpopulations compared to snRNA-seq, with satglia and fibroblasts remaining among the top differentially regulated populations. In OSM-treated satglia (Fig. 8F), upregulated genes included *CCL2*, promoting immune recruitment; *NRG3* and *MAML2*, regulating neural-glial signaling; and *MACF1* and *FNDC3A*, supporting cytoskeletal integrity. *DYRK1A*, *FTO*, and *KMT2C* reflected signaling and metabolic shifts, while *SMAD1*, *TMEM132A*, and *THBS2*, and *MMP14* driving stress responses, and ECM remodeling. *VPS13C*, *SEL1L*, and *AAK1* underscored vesicle trafficking and protein degradation, collectively defining satglia response to OSM. In OSM-treated Fibroblasts (Fig. 8G), upregulated genes include *PIEZO1*, *COL23A1*, and *MMP15*, which drive mechanotransduction and ECM remodeling.

**Figure 8.**
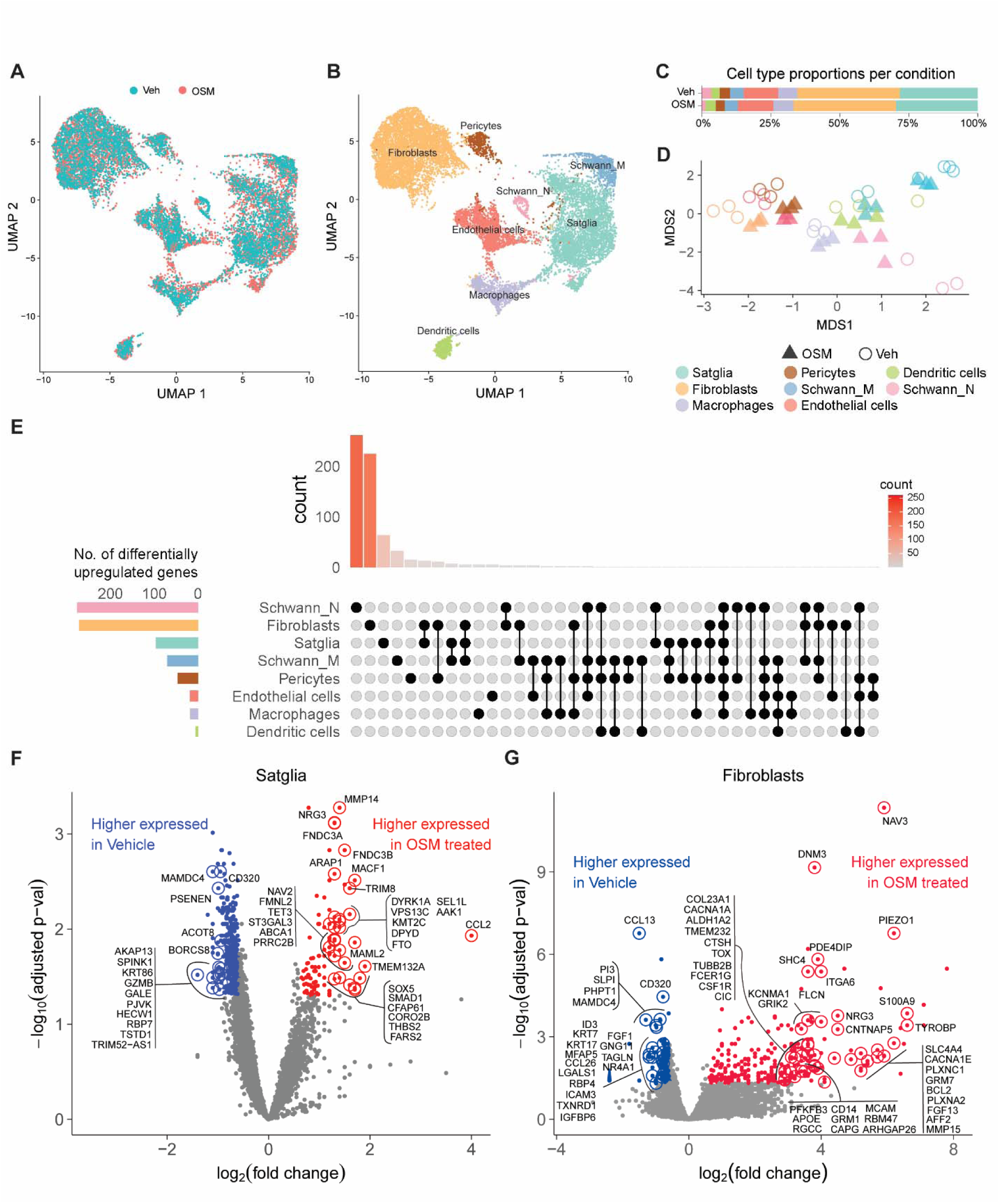
OSM induces distinct transcriptional changes across multiple cell populations in scRNAseq analysis of hDRG cultures. **A**) UMAP projection showing overlapping distribution in the vehicle and OSM-treated conditions. DIV4 hDRG cultures were treated with 10 ng/mL OSM for 6 hours. N = 3 replicates per condition. **B**) UMAP plot of annotated cell types, including Schwann cells (Schwann_M and Schwann_N), satglia, fibroblasts, macrophages, dendritic cells, endothelial cells, and pericytes. **C**) Bar plot depicting the proportion of each cell type in the vehicle and OSM-treated conditions. **D**) MDS plot of sample clustering by cell type. **E**) Upset plot showing the number of differentially upregulated genes per cell type in OSM treatment compared to the vehicle, with the highest count in Schwann_N cells, fibroblasts, and satglia. **F**) Volcano plot of differentially expressed genes in satglia, with genes higher expressed in OSM-treated (red) compared to vehicle (blue). Log2FC ≥ 0.585, adjusted p-value < 0.05 for upregulated genes and Log2FC ≤ -0.585 for downregulated genes. **G**) Volcano plot of differentially expressed genes in fibroblasts, similarly contrasting higher expression in OSM versus vehicle-treated conditions.

To gain a more comprehensive understanding of the effects of OSM at both the mRNA and protein levels within DRGs, we sought to investigate changes in the proteomic landscape. Proteomic analysis was performed on acutely sliced hDRG explants (4 Vehicle and 4 OSM) using the SomaScan platform (Fig. S9A and B). Our analysis revealed 220 upregulated proteins, including those encoded by genes such as *ICAM1*, *FMR1*, *IGF1R*, *PTPN9*, *TLR1*, *TLR3*, *COL6A3*, *SEMA3E*, *SERPINH1*, *TRIM28*, *MAPK13*, *EEF1A1*, and *EIF2AK4*, (Fig. S9B and C). Many of the upregulated proteins are involved in the TNF, VEGF, and P13K-Akt signaling pathways, which are also implicated in pain. Finally, to integrate our findings, we performed an analysis on the OSM-treated proteomics data, as well as bulk RNAseq, snRNAseq, and scRNAseq data (Fig. S10). While we observed few correlations between the OSM proteomics data and both the OSM bulk RNAseq and pain bulk RNAseq datasets, we noted that some of the most robustly upregulated proteins correlated with increases in gene expression in specific cell types, most notably in the OSM scRNAseq data (Fig. S10). Altogether, our multi-omics approach provides clear evidence that OSM can drive distinct molecular responses, both at the transcript and protein level, across multiple cell populations in the hDRG.

## DISCUSSION

Our findings demonstrate that OSM-OSMR signaling is an excellent candidate for orchestrating molecular and physiological changes in the hDRG that cause at least some aspects of neuropathic pain in patients. First, OSMR is expressed by multiple subsets of nociceptive DRG neurons in humans, as well as satglia, macrophages, and likely fibroblasts. OSM is found in macrophages within the DRG making these cells a prime candidate for the cell type that might release OSM in the context of neuropathic pain. The cytokine activates MAPK signaling in hDRG neurons, exemplified by MNK1/2 activation and eIF4E phosphorylation. This arm of the MAPK signaling pathway is linked to neuropathic pain in animal models (*65, 67, 69, 88, 89*), and inhibition of MNK signaling reduces SA in hDRG neurons recovered from individuals with neuropathic pain (*8*). Consistent with this, OSM causes action potential firing in hDRG neurons from organ donors with no history of neuropathic pain. The lasting effect of OSM on these neurons is reversed when MNK activity is inhibited with the specific inhibitor eFT508.

Finally, OSM treatment of hDRG cultures causes transcriptomic changes that closely resemble those observed in patients with neuropathic pain (*7, 9*). Altogether, these findings illustrate that OSM signaling is sufficient to drive a state in hDRG that recapitulates key molecular and physiological findings in neuropathic pain patients.

Genetic (*14*), basic science clinical (*15, 17*) and pre-clinical studies (*17*), and clinical trial data (*16*) support the hypothesis that OSM is involved in skin diseases that cause itch in patients. Although some previous animal studies support a role for OSM in pain (*27–29*), why have these clinical studies failed to link the cytokine to pain in humans? One key factor is that the OSMR mutations linked to itch in FPLCA are a loss of function mutation that renders the receptor incapable of transducing signals from either OSM or IL31 (*14*). Loss of function of OSMR in FPLCA seems to cause changes in keratinocyte cell cycle leading to bumpy keratinous deposits in skin that then become itchy, causing this symptom in the disease. Other pruritic diseases like prurigo nodularis are associated with OSM upregulation in skin, and patients experience both itch and pain (*15, 90*). Interestingly, pain in prurigo nodularis is often described in terms that are associated with neuropathic pain such as burning and shooting pain and is often treated with gabapentinoids (*90*). Vixarelimab, which blocks OSMR signaling by OSM and IL31, effectively reduces itching and skin lesions in prurigo nodularis patients, but any effects on pain were not reported in that study and there were no pain endpoints in the published trial protocol (*16*). We provide clear evidence that OSMR is expressed in multiple classes of human nociceptors, all of which also express the signal transducer gp130 (*54, 91*), and that these neurons innervate the outmost layers of skin, where itch signals are produced, and deeper layers, where pain signals from the skin originate. We anticipate that further examination of symptom resolution in pruritic disease trials for vixarelimab or similar therapeutics will find positive effects of these biologics/drugs on pain in these patients.

We describe a profound species difference in OSMR expression between mice and humans wherein there is an expansion of expression in humans to multiple classes of nociceptors from the relatively specific expression of OSMR in pruriceptive neurons in adult mice (*17, 59*). What drives this difference in expression across species? While the precise mechanisms are not currently known, we speculate that this may be functionally linked to an expansion of the TrkA-positive subset of neurons in the hDRG (*54, 92–94*). In mice, TrkA is expressed in all DRG neurons in early life but is shut off after birth in neurons that become the so-called non-peptidergic class of nociceptors (*95*). Previous studies show that OSM is required for the development of a large subset of mostly non-peptidergic nociceptors wherein OSM knockout mice lose ∼ 20-30% of nociceptors and have decreased nociceptive responses to heat and chemical irritants (*27*). In humans, this distinction between peptidergic and non-peptidergic nociceptors appears to be lost because all human nociceptors express the peptidergic markers calcitonin gene-related peptide (CGRP) and TrkA and all human nociceptors also express TRPV1 (*92, 93*). This fundamental difference supports the idea that transcriptional programs controlling gene expression in nociceptors are under selection pressure, and *OSMR* is likely one of these genes. Genomic-based structural differences in OSM that change the pharmacology of the cytokine across species also support this conclusion for the cytokine-receptor signaling module (*22*). While previous studies have examined how genomic variation in TRP channels play a role in climatic adaptation for animals like ground squirrels and camels, or for capsaicin sensitivity in birds (*96, 97*), little attention has been paid to evolutionary adaptation that may contribute to neuro-immune interactions that make certain species, like humans, susceptible to neuropathic pain.

## LIMITATIONS

A significant limitation is that our work on hDRG can only give us insight into regulation of the activity of the peripheral cells we have studied in this work. Ultimately pain is a sensory and emotional experience that requires brain processing, which is beyond the experimental scope of our work. Activity in DRG neurons is a key driver of neuropathic pain as exemplified by many clinical studies that demonstrate a rapid alleviation of pain with peripheral nerve block in neuropathic pain patients (*4–6*). We hypothesize that patients with a clear peripheral driver of their neuropathic pain are the most likely to potentially benefit from OSM/OSMR-targeting therapeutics.

OSM/OSMR signaling is unlikely to act alone to produce neuropathic pain. Instead, OSM upregulation in the DRG or in painful tissues should be viewed as part of a network of changes that drives the disease state. From this perspective, our work is limited by its reductionist approach to studying OSM in isolation. RNA sequencing studies on hDRG samples from patients suffering from neuropathic pain show transcriptional changes that are likely driven by immune cell state alterations within the DRG (*7, 9, 10*). Similar changes may also occur in the peripheral nerve and in peripheral tissues. Single cell and/or spatial transcriptomic studies can give deeper insight into how these changes in specific immune cells drive signaling networks in the neuro-immune axis to cause pain. Recent studies using single nucleus sequencing from synovial tissues in treatment resistant rheumatoid arthritis (*21*) and in chronic low back pain (*98*) show the power of this approach for identifying patient subtypes based on cell-states and how this can be linked to clinical phenotypes, like pain. Increased access to hDRG tissues accompanied by detailed medical histories can enable recovery of tissues that can make these studies possible (*99*). Ultimately these studies will likely be needed to understand how to target OSM/OSMR signaling for the treatment of pain. While clinical trials in neuropathic pain for biologics like vixarelimab are warranted based on our findings, we envision approaches that alter pathological immune cell states as an important future approach to treating neuropathic pain. Such an approach has been used in preclinical pain models with success (*100*).

Another of our study’s limitations is the underrepresentation or depletion of neuronal clusters following the cell capture techniques used for scRNAseq and snRNAseq. This constrained our ability to fully investigate neuron-specific transcriptional changes in the hDRGs following OSM exposure. To address this challenge, future studies could use spatial transcriptomics and laser capture microdissection for more precise capture of neuronal populations, allowing for a deeper analysis of neuronal gene expression and spatial organization within the DRG. Nevertheless, the combination of cellular signaling and electrophysiological recordings couples with transcriptomic studies on OSM signaling in the DRG shows that this cytokine can have profound effects on multiple subtypes of neurons, glia and other cell types in the hDRG.

## CONCLUSION

Our work is an example of the importance of a human-first approach to understanding neuropathic pain mechanisms. Prior to our experiments, major species differences in OSM/OSMR signaling were identified informing our hypotheses leading to these studies (*23–25*). We discovered pervasive differences in OSMR localization and signaling in hDRG where OSM leads to direct excitation of hDRG neurons through MAPK signaling and induces transcriptional changes that mimic those seen in neuropathic pain patients. OSM/OSMR-signaling is a bona-fide target for the treatment of neuropathic pain.

## Acknowledgements

The authors thank the organ donors and their families for their gift of life.

Conflict of Interest Statement

T.J.P. is a co-founder and holds equity in 4E Therapeutics, NuvoNuro, PARMedics, Nerveli. T.J.P. has received research grants from AbbVie, Eli Lilly, Grunenthal, Evommune, Hoba Therapeutics, and The National Institutes of Health. M.S.Y. is a co-founder and holds equity in NuvoNuro. M.C. is chief medical officer and holds options in 4E Therapeutics.

## REFERENCES CITED

1. N. B. Finnerup, N. Attal, S. Haroutounian, E. McNicol, R. Baron, R. H. Dworkin, I. Gilron, M. Haanpaa, P. Hansson, T. S. Jensen, P. R. Kamerman, K. Lund, A. Moore, S. N. Raja, A. S. Rice, M. Rowbotham, E. Sena, P. Siddall, B. H. Smith, M. Wallace, Pharmacotherapy for neuropathic pain in adults: a systematic review and meta-analysis. Lancet Neurol 14, 162–173 (2015).

2. S. J. Middleton, A. M. Barry, M. Comini, Y. Li, P. R. Ray, S. Shiers, A. C. Themistocleous, M. L. Uhelski, X. Yang, P. M. Dougherty, T. J. Price, D. L. Bennett, Studying human nociceptors: from fundamentals to clinic. Brain 144, 1312–1335 (2021).

3. K. E. Sadler, J. S. Mogil, C. L. Stucky, Innovations and advances in modelling and measuring pain in animals. Nat Rev Neurosci 23, 70–85 (2022).

4. A. Vaso, H. M. Adahan, A. Gjika, S. Zahaj, T. Zhurda, G. Vyshka, M. Devor, Peripheral nervous system origin of phantom limb pain. Pain 155, 1384–1391 (2014).

5. S. Haroutounian, A. L. Ford, K. Frey, L. Nikolajsen, N. B. Finnerup, A. Neiner, E. D. Kharasch, P. Karlsson, M. M. Bottros, How central is central poststroke pain? The role of afferent input in poststroke neuropathic pain: a prospective, open-label pilot study. Pain, (2018).

6. S. Haroutounian, L. Nikolajsen, T. F. Bendtsen, N. B. Finnerup, A. D. Kristensen, J. B. Hasselstrom, T. S. Jensen, Primary afferent input critical for maintaining spontaneous pain in peripheral neuropathy. Pain 155, 1272–1279 (2014).

7. R. Y. North, Y. Li, P. Ray, L. D. Rhines, C. E. Tatsui, G. Rao, C. A. Johansson, H. Zhang, Y. H. Kim, B. Zhang, G. Dussor, T. H. Kim, T. J. Price, P. M. Dougherty, Electrophysiological and transcriptomic correlates of neuropathic pain in human dorsal root ganglion neurons. Brain 142, 1215–1226 (2019).

8. Y. Li, M. L. Uhelski, R. Y. North, J. M. Mwirigi, C. E. Tatsui, J. P. Cata, G. Corrales, T. J. Price, P. M. Dougherty, MNK inhibitor eFT508 (Tomivosertib) suppresses ectopic activity in human dorsal root ganglion neurons from dermatomes with radicular neuropathic pain. bioRxiv, (2023).

9. P. R. Ray, S. Shiers, J. P. Caruso, D. Tavares-Ferreira, I. Sankaranarayanan, M. L. Uhelski, Y. Li, R. Y. North, C. Tatsui, G. Dussor, M. D. Burton, P. M. Dougherty, T. J. Price, RNA profiling of human dorsal root ganglia reveals sex differences in mechanisms promoting neuropathic pain. Brain 146, 749–766 (2023).

10. B. E. Hall, E. Macdonald, M. Cassidy, S. Yun, M. R. Sapio, P. Ray, M. Doty, P. Nara, M. D. Burton, S. Shiers, A. Ray-Chaudhury, A. J. Mannes, T. J. Price, M. J. Iadarola, A. B. Kulkarni, Transcriptomic analysis of human sensory neurons in painful diabetic neuropathy reveals inflammation and neuronal loss. Sci Rep 12, 4729 (2022).

11. S. Rosen, B. Ham, J. S. Mogil, Sex differences in neuroimmunity and pain. J Neurosci Res 95, 500–508 (2017).

12. J. S. Mogil, A. L. Bailey, Sex and gender differences in pain and analgesia. Prog Brain Res 186, 141–157 (2010).

13. C. L. Wolf, C. Pruett, D. Lighter, C. L. Jorcyk, The clinical relevance of OSM in inflammatory diseases: a comprehensive review. Front Immunol 14, 1239732 (2023).

14. K. Arita, A. P. South, G. Hans-Filho, T. H. Sakuma, J. Lai-Cheong, S. Clements, M. Odashiro, D. N. Odashiro, G. Hans-Neto, N. R. Hans, M. V. Holder, B. S. Bhogal, S. T. Hartshorne, M. Akiyama, H. Shimizu, J. A. McGrath, Oncostatin M receptor-beta mutations underlie familial primary localized cutaneous amyloidosis. Am J Hum Genet 82, 73–80 (2008).

15. T. Hashimoto, L. A. Nattkemper, H. S. Kim, C. D. Kursewicz, E. Fowler, S. M. Shah, S. Nanda, R. A. Fayne, J. F. Paolini, P. Romanelli, G. Yosipovitch, Itch intensity in prurigo nodularis is closely related to dermal interleukin-31, oncostatin M, IL-31 receptor alpha and oncostatin M receptor beta. Exp Dermatol 30, 804–810 (2021).

16. H. Sofen, R. Bissonnette, G. Yosipovitch, J. I. Silverberg, S. Tyring, W. J. Loo, M. Zook, M. Lee, L. Zou, G. L. Jiang, J. F. Paolini, Efficacy and safety of vixarelimab, a human monoclonal oncostatin M receptor beta antibody, in moderate-to-severe prurigo nodularis: a randomised, double-blind, placebo-controlled, phase 2a study. EClinicalMedicine 57, 101826 (2023).

17. P. Y. Tseng, M. A. Hoon, Oncostatin M can sensitize sensory neurons in inflammatory pruritus. Sci Transl Med 13, eabe3037 (2021).

18. W. Chen, Y. Li, M. Steinhoff, W. Zhang, J. Buddenkotte, T. Buhl, R. Zhu, X. Yan, Z. Lu, S. Xiao, J. Wang, J. Meng, The PLAUR signaling promotes chronic pruritus. FASEB J 36, e22368 (2022).

19. J. Deng, V. Parthasarathy, M. Marani, Z. Bordeaux, K. Lee, C. Trinh, H. L. Cornman, A. Kambala, T. Pritchard, S. Chen, N. Sutaria, O. O. Oladipo, M. M. Kwatra, M. P. Alphonse, S. G. Kwatra, Extracellular matrix and dermal nerve growth factor dysregulation in prurigo nodularis compared to atopic dermatitis. Front Med (Lausanne) 9, 1022889 (2022).

20. H. M. Hermanns, Oncostatin M and interleukin-31: Cytokines, receptors, signal transduction and physiology. Cytokine Growth Factor Rev 26, 545–558 (2015).

21. F. Zhang, A. H. Jonsson, A. Nathan, N. Millard, M. Curtis, Q. Xiao, M. Gutierrez-Arcelus, W. Apruzzese, G. F. M. Watts, D. Weisenfeld, S. Nayar, J. Rangel-Moreno, N. Meednu, K. E. Marks, I. Mantel, J. B. Kang, L. Rumker, J. Mears, K. Slowikowski, K. Weinand, D. E. Orange, L. Geraldino-Pardilla, K. D. Deane, D. Tabechian, A. Ceponis, G. S. Firestein, M. Maybury, I. Sahbudin, A. Ben-Artzi, A. M. Mandelin, 2nd, A. Nerviani, M. J. Lewis, F. Rivellese, C. Pitzalis, L. B. Hughes, D. Horowitz, E. DiCarlo, E. M. Gravallese, B. F. Boyce, R. A. S. L. E. N. Accelerating Medicines Partnership, L. W. Moreland, S. M. Goodman, H. Perlman, V. M. Holers, K. P. Liao, A. Filer, V. P. Bykerk, K. Wei, D. A. Rao, L. T. Donlin, J. H. Anolik, M. B. Brenner, S. Raychaudhuri, Deconstruction of rheumatoid arthritis synovium defines inflammatory subtypes. Nature 623, 616–624 (2023).

22. J. M. Adrian-Segarra, K. Sreenivasan, P. Gajawada, H. Lorchner, T. Braun, J. Poling, The AB loop of oncostatin M (OSM) determines species-specific signaling in humans and mice. J Biol Chem 293, 20181–20199 (2018).

23. R. A. Lindberg, T. S. Juan, A. A. Welcher, Y. Sun, R. Cupples, B. Guthrie, F. A. Fletcher, Cloning and characterization of a specific receptor for mouse oncostatin M. Mol Cell Biol 18, 3357–3367 (1998).

24. J. Drechsler, J. Grotzinger, H. M. Hermanns, Characterization of the rat oncostatin M receptor complex which resembles the human, but differs from the murine cytokine receptor. PLoS One 7, e43155 (2012).

25. T. Hara, M. Ichihara, A. Yoshimura, A. Miyajima, Cloning and biological activity of murine oncostatin M. Leukemia 11 Suppl 3, 449–450 (1997).

26. M. Suehiro, T. Numata, R. Saito, N. Yanagida, C. Ishikawa, K. Uchida, T. Kawaguchi, Y. Yanase, Y. Ishiuji, J. McGrath, A. Tanaka, Oncostatin M suppresses IL31RA expression in dorsal root ganglia and interleukin-31-induced itching. Front Immunol 14, 1251031 (2023).

27. Y. Morikawa, S. Tamura, K. Minehata, P. J. Donovan, A. Miyajima, E. Senba, Essential function of oncostatin m in nociceptive neurons of dorsal root ganglia. J Neurosci 24, 1941–1947 (2004).

28. M. Langeslag, C. E. Constantin, M. Andratsch, S. Quarta, N. Mair, M. Kress, Oncostatin M induces heat hypersensitivity by gp130-dependent sensitization of TRPV1 in sensory neurons. Mol Pain 7, 102 (2011).

29. A. Garza Carbajal, A. Ebersberger, A. Thiel, L. Ferrari, J. Acuna, S. Brosig, J. Isensee, K. Moeller, M. Siobal, S. Rose-John, J. Levine, H. G. Schaible, T. Hucho, Oncostatin M induces hyperalgesic priming and amplifies signaling of cAMP to ERK by RapGEF2 and PKA. J Neurochem 157, 1821–1837 (2021).

30. J. Koerner, D. Howard, H. J. Solinski, M. M. Moreno, N. Haag, A. Fiebig, S. A. Bhuiyan, I. Toklucu, R. Bott, I. Sankaranarayanan, D. Tavares-Ferreira, S. Shiers, N. N. Inturi, A. Maxion, L. Ernst, L. Bonaguro, J. Schulte-Schrepping, M. D. Beyer, T. Stiehl, W. Renthal, I. Kurth, T. Price, M. Schmelz, B. Namer, S. Tripathy, A. Lampert, Molecular architecture of human dermal sleeping nociceptors. bioRxiv, 2024.2012.2020.629638 (2024).

31. Y. Li, M. L. Uhelski, R. Y. North, J. M. Mwirigi, C. E. Tatsui, K. E. McDonough, J. P. Cata, G. Corrales, G. Dussor, T. J. Price, P. M. Dougherty, Tomivosertib reduces ectopic activity in dorsal root ganglion neurons from patients with radiculopathy. Brain 147, 2991–2997 (2024).

32. S. Shiers, J. J. Sahn, T. J. Price, MNK1 and MNK2 Expression in the Human Dorsal Root and Trigeminal Ganglion. Neuroscience 515, 96–107 (2023).

33. S. Shiers, R. M. Klein, T. J. Price, Quantitative differences in neuronal subpopulations between mouse and human dorsal root ganglia demonstrated with RNAscope in situ hybridization. Pain, (2020).

34. C. Stringer, T. Wang, M. Michaelos, M. Pachitariu, Cellpose: a generalist algorithm for cellular segmentation. Nat Methods 18, 100–106 (2021).

35. R. Ali Marandi Ghoddousi, V. M. Magalong, A. K. Kamitakahara, P. Levitt, SCAMPR, a single-cell automated multiplex pipeline for RNA quantification and spatial mapping. Cell Rep Methods 2, 100316 (2022).

36. U. Franco-Enzastiga, N. N. Inturi, K. Natarajan, J. M. Mwirigi, K. Mazhar, J. C. M. Schlachetzki, M. Schumacher, T. J. Price, Epigenomic landscape of the human dorsal root ganglion: sex differences and transcriptional regulation of nociceptive genes. Pain 166, 614–630 (2025).

37. A. Butler, P. Hoffman, P. Smibert, E. Papalexi, R. Satija, Integrating single-cell transcriptomic data across different conditions, technologies, and species. Nat Biotechnol 36, 411–420 (2018).

38. T. Stuart, A. Butler, P. Hoffman, C. Hafemeister, E. Papalexi, W. M. Mauck, 3rd, Y. Hao, M. Stoeckius, P. Smibert, R. Satija, Comprehensive Integration of Single-Cell Data. Cell 177, 1888–1902 e1821 (2019).

39. P. J. Albrecht, G. Houk, E. Ruggiero, M. Dockum, M. Czerwinski, J. Betts, J. P. Wymer, C. E. Argoff, F. L. Rice, Keratinocyte Biomarkers Distinguish Painful Diabetic Peripheral Neuropathy Patients and Correlate With Topical Lidocaine Responsiveness. Front Pain Res (Lausanne) 2, 790524 (2021).

40. D. Bowsher, C. Geoffrey Woods, A. K. Nicholas, O. M. Carvalho, C. E. Haggett, B. Tedman, J. M. Mackenzie, D. Crooks, N. Mahmood, J. A. Twomey, S. Hann, D. Jones, J. P. Wymer, P. J. Albrecht, C. E. Argoff, F. L. Rice, Absence of pain with hyperhidrosis: a new syndrome where vascular afferents may mediate cutaneous sensation. Pain 147, 287–298 (2009).

41. P. J. Albrecht, S. Hines, E. Eisenberg, D. Pud, D. R. Finlay, K. M. Connolly, M. Pare, G. Davar, F. L. Rice, Pathologic alterations of cutaneous innervation and vasculature in affected limbs from patients with complex regional pain syndrome. Pain 120, 244–266 (2006).

42. M. Fetell, M. Sendel, T. Li, L. Marinelli, J. Vollert, E. Ruggerio, G. Houk, M. Dockum, P. J. Albrecht, F. L. Rice, R. Baron, Cutaneous nerve fiber and peripheral Nav1.7 assessment in a large cohort of patients with postherpetic neuralgia. Pain 164, 2435–2446 (2023).

43. F. L. Rice, G. Houk, J. P. Wymer, S. J. C. Gosline, J. Guinney, J. Wu, N. Ratner, M. P. Jankowski, S. La Rosa, M. Dockum, J. R. Storey, S. L. Carroll, P. J. Albrecht, V. M. Riccardi, The evolution and multi-molecular properties of NF1 cutaneous neurofibromas originating from C-fiber sensory endings and terminal Schwann cells at normal sites of sensory terminations in the skin. PLoS One 14, e0216527 (2019).

44. P. J. Albrecht, Q. Hou, C. E. Argoff, J. R. Storey, J. P. Wymer, F. L. Rice, Excessive peptidergic sensory innervation of cutaneous arteriole-venule shunts (AVS) in the palmar glabrous skin of fibromyalgia patients: implications for widespread deep tissue pain and fatigue. Pain Med 14, 895–915 (2013).

45. F. L. Rice, B. T. Fundin, J. Arvidsson, H. Aldskogius, O. Johansson, Comprehensive immunofluorescence and lectin binding analysis of vibrissal follicle sinus complex innervation in the mystacial pad of the rat. J Comp Neurol 385, 149–184 (1997).

46. S. Palomino, K. Gabriel, J. Mwirigi, A. Cervantes, P. Horton, G. Funk, A. Moutal, L. Martin, R. Khanna, T. Price, A. Patwardhan, Genetic editing of primary human dorsal root ganglion neurons using CRISPR-Cas9 with functional confirmation. bioRxiv, 2024.2004.2002.587857 (2024).

47. J. M. M. I. Sankaranarayanan, D. Tavares Ferreira, and T. J. Price. (SPARC, 2024).

48. M. J. Lacagnina, K. F. Willcox, N. Boukelmoune, A. Bavencoffe, I. Sankaranarayanan, D. T. Barratt, Y. A. Zuberi, D. Dayani, M. V. Chavez, J. T. Lu, A. B. Farinotti, S. Shiers, A. M. Barry, J. M. Mwirigi, D. Tavares-Ferreira, G. A. Funk, A. M. Cervantes, C. I. Svensson, E. T. Walters, M. R. Hutchinson, C. J. Heijnen, T. J. Price, N. T. Fiore, P. M. Grace, B cells drive neuropathic pain-related behaviors in mice through IgG-Fc gamma receptor signaling. Sci Transl Med 16, eadj1277 (2024).

49. M. D. Young, S. Behjati, SoupX removes ambient RNA contamination from droplet-based single-cell RNA sequencing data. Gigascience 9, (2020).

50. J. Lackovic, T. J. Price, G. Dussor, MNK1/2 contributes to periorbital hypersensitivity and hyperalgesic priming in preclinical migraine models. Brain 146, 448–454 (2023).

51. S. A. Bhuiyan, M. Xu, L. Yang, E. Semizoglou, P. Bhatia, K. I. Pantaleo, I. Tochitsky, A. Jain, B. Erdogan, S. Blair, V. Cat, J. M. Mwirigi, I. Sankaranarayanan, D. Tavares-Ferreira, U. Green, L. A. McIlvried, B. A. Copits, Z. Bertels, J. S. Del Rosario, A. J. Widman, R. A. Slivicki, J. Yi, R. Sharif-Naeini, C. J. Woolf, J. K. Lennerz, J. L. Whited, T. J. Price, W. G. I. Robert, W. Renthal, Harmonized cross-species cell atlases of trigeminal and dorsal root ganglia. Sci Adv 10, eadj9173 (2024).

52. H. L. Crowell, C. Soneson, P. L. Germain, D. Calini, L. Collin, C. Raposo, D. Malhotra, M. D. Robinson, muscat detects subpopulation-specific state transitions from multi-sample multi-condition single-cell transcriptomics data. Nat Commun 11, 6077 (2020).

53. D. T. Ferreira, B. Q. Shen, J. M. Mwirigi, S. Shiers, I. Sankaranarayanan, M. Kotamarti, N. N. Inturi, K. Mazhar, E. E. Ubogu, G. Thomas, T. Lalli, D. Wukich, T. J. Price, Deciphering the molecular landscape of human peripheral nerves: implications for diabetic peripheral neuropathy. bioRxiv, (2024).

54. D. Tavares-Ferreira, S. Shiers, P. R. Ray, A. Wangzhou, V. Jeevakumar, I. Sankaranarayanan, A. M. Cervantes, J. C. Reese, A. Chamessian, B. A. Copits, P. M. Dougherty, R. W. T. Gereau, M. D. Burton, G. Dussor, T. J. Price, Spatial transcriptomics of dorsal root ganglia identifies molecular signatures of human nociceptors. Sci Transl Med 14, eabj8186 (2022).

55. V. Prato, F. J. Taberner, J. R. F. Hockley, G. Callejo, A. Arcourt, B. Tazir, L. Hammer, P. Schad, P. A. Heppenstall, E. S. Smith, S. G. Lechner, Functional and Molecular Characterization of Mechanoinsensitive "Silent" Nociceptors. Cell Rep 21, 3102–3115 (2017).

56. O. Avraham, R. Feng, E. E. Ewan, J. Rustenhoven, G. Zhao, V. Cavalli, Profiling sensory neuron microenvironment after peripheral and central axon injury reveals key pathways for neural repair. Elife 10, (2021).

57. A. A. Mapps, M. B. Thomsen, E. Boehm, H. Zhao, S. Hattar, R. Kuruvilla, Diversity of satellite glia in sympathetic and sensory ganglia. Cell Rep 38, 110328 (2022).

58. U. Franco-Enzastiga, N. N. Inturi, K. Natarajan, J. M. Mwirigi, K. Mazhar, J. C. M. Schlachetzki, M. Schumacher, T. J. Price, Epigenomic landscape of the human dorsal root ganglion: sex differences and transcriptional regulation of nociceptive genes. bioRxiv, (2024).

59. S. A. Bhuiyan, M. Xu, L. Yang, E. Semizoglou, P. Bhatia, K. I. Pantaleo, I. Tochitsky, A. Jain, B. Erdogan, S. Blair, V. Cat, J. M. Mwirigi, I. Sankaranarayanan, D. Tavares-Ferreira, U. Green, L. A. McIlvried, B. A. Copits, Z. Bertels, J. S. Del Rosario, A. J. Widman, R. A. Slivicki, J. Yi, R. Sharif-Naeini, C. J. Woolf, J. K. Lennerz, J. L. Whited, T. J. Price, W. Renthal, Harmonized cross-species cell atlases of trigeminal and dorsal root ganglia. Science Advances 10, eadj9173 (2024).

60. J. Körner, D. Howard, H. J. Solinski, M. M. Moreno, N. Haag, A. Fiebig, S. A. Bhuiyan, I. Toklucu, R. Bott, I. Sankaranarayanan, D. Tavares-Ferreira, S. Shiers, N. N. Inturi, A. Maxion, L. Ernst, L. Bonaguro, J. Schulte-Schrepping, M. D. Beyer, T. Stiehl, W. Renthal, I. Kurth, T. Price, M. Schmelz, B. Namer, S. Tripathy, A. Lampert, Molecular architecture of human dermal sleeping nociceptors. bioRxiv, 2024.2012.2020.629638 (2024).

61. M. J. Zylka, F. L. Rice, D. J. Anderson, Topographically distinct epidermal nociceptive circuits revealed by axonal tracers targeted to Mrgprd. Neuron 45, 17–25 (2005).

62. M. Pare, P. J. Albrecht, C. J. Noto, N. L. Bodkin, G. L. Pittenger, D. J. Schreyer, X. T. Tigno, B. C. Hansen, F. L. Rice, Differential hypertrophy and atrophy among all types of cutaneous innervation in the glabrous skin of the monkey hand during aging and naturally occurring type 2 diabetes. J Comp Neurol 501, 543–567 (2007).

63. L. Dollet, L. S. Lundell, A. V. Chibalin, L. A. Pendergrast, N. J. Pillon, E. L. Lansbury, M. Elmastas, S. Frendo-Cumbo, J. Jalkanen, T. de Castro Barbosa, D. T. Cervone, K. Caidahl, O. Dmytriyeva, A. S. Deshmukh, R. Barres, M. Ryden, H. Wallberg-Henriksson, J. R. Zierath, A. Krook, Exercise-induced crosstalk between immune cells and adipocytes in humans: Role of oncostatin-M. Cell Rep Med 5, 101348 (2024).

64. N. R. West, A. N. Hegazy, B. M. J. Owens, S. J. Bullers, B. Linggi, S. Buonocore, M. Coccia, D. Gortz, S. This, K. Stockenhuber, J. Pott, M. Friedrich, G. Ryzhakov, F. Baribaud, C. Brodmerkel, C. Cieluch, N. Rahman, G. Muller-Newen, R. J. Owens, A. A. Kuhl, K. J. Maloy, S. E. Plevy, I. B. D. C. I. Oxford, S. Keshav, S. P. L. Travis, F. Powrie, Oncostatin M drives intestinal inflammation and predicts response to tumor necrosis factor-neutralizing therapy in patients with inflammatory bowel disease. Nat Med 23, 579–589 (2017).

65. J. K. Moy, A. Khoutorsky, M. N. Asiedu, B. J. Black, J. L. Kuhn, P. Barragan-Iglesias, S. Megat, M. D. Burton, C. C. Burgos-Vega, O. K. Melemedjian, S. Boitano, J. Vagner, C. G. Gkogkas, J. J. Pancrazio, J. S. Mogil, G. Dussor, N. Sonenberg, T. J. Price, The MNK-eIF4E Signaling Axis Contributes to Injury-Induced Nociceptive Plasticity and the Development of Chronic Pain. J Neurosci 37, 7481–7499 (2017).

66. M. E. Mitchell, G. Torrijos, L. F. Cook, J. M. Mwirigi, L. He, S. Shiers, T. J. Price, Interleukin-6 induces nascent protein synthesis in human dorsal root ganglion nociceptors primarily via MNK-eIF4E signaling. Neurobiol Pain 16, 100159 (2024).

67. S. Shiers, J. Mwirigi, G. Pradhan, M. Kume, B. Black, P. Barragan-Iglesias, J. K. Moy, G. Dussor, J. J. Pancrazio, S. Kroener, T. J. Price, Reversal of peripheral nerve injury-induced neuropathic pain and cognitive dysfunction via genetic and tomivosertib targeting of MNK. Neuropsychopharmacology 45, 524–533 (2020).

68. J. K. Moy, J. L. Kuhn, T. A. Szabo-Pardi, G. Pradhan, T. J. Price, eIF4E phosphorylation regulates ongoing pain, independently of inflammation, and hyperalgesic priming in the mouse CFA model. Neurobiol Pain 4, 45–50 (2018).

69. S. Megat, P. R. Ray, J. K. Moy, T. F. Lou, P. Barragan-Iglesias, Y. Li, G. Pradhan, A. Wanghzou, A. Ahmad, M. D. Burton, R. Y. North, P. M. Dougherty, A. Khoutorsky, N. Sonenberg, K. R. Webster, G. Dussor, Z. T. Campbell, T. J. Price, Nociceptor Translational Profiling Reveals the Ragulator-Rag GTPase Complex as a Critical Generator of Neuropathic Pain. J Neurosci 39, 393–411 (2019).

70. P. Barragan-Iglesias, U. Franco-Enzastiga, V. Jeevakumar, S. Shiers, A. Wangzhou, V. Granados-Soto, Z. T. Campbell, G. Dussor, T. J. Price, Type I Interferons Act Directly on Nociceptors to Produce Pain Sensitization: Implications for Viral Infection-Induced Pain. J Neurosci 40, 3517–3532 (2020).

71. A. Bavencoffe, E. A. Spence, M. Y. Zhu, A. Garza-Carbajal, K. E. Chu, O. E. Bloom, C. W. Dessauer, E. T. Walters, Macrophage Migration Inhibitory Factor (MIF) Makes Complex Contributions to Pain-Related Hyperactivity of Nociceptors after Spinal Cord Injury. J Neurosci 42, 5463–5480 (2022).

72. A. G. Bavencoffe, E. R. Lopez, K. N. Johnson, J. Tian, F. M. Gorgun, B. Q. Shen, M. X. Zhu, C. W. Dessauer, E. T. Walters, Widespread latent hyperactivity of nociceptors outlasts enhanced avoidance behavior following incision injury. bioRxiv, (2024).

73. S. C. Berkey, J. J. Herrera, M. A. Odem, S. Rahman, S. S. Cheruvu, X. Cheng, E. T. Walters, C. W. Dessauer, A. G. Bavencoffe, EPAC1 and EPAC2 promote nociceptor hyperactivity associated with chronic pain after spinal cord injury. Neurobiol Pain 7, 100040 (2020).

74. G. Laumet, A. Bavencoffe, J. D. Edralin, X. J. Huo, E. T. Walters, R. Dantzer, C. J. Heijnen, A. Kavelaars, Interleukin-10 resolves pain hypersensitivity induced by cisplatin by reversing sensory neuron hyperexcitability. Pain 161, 2344–2352 (2020).

75. E. R. Lopez, A. G. Carbajal, J. B. Tian, A. Bavencoffe, M. X. Zhu, C. W. Dessauer, E. T. Walters, Serotonin enhances depolarizing spontaneous fluctuations, excitability, and ongoing activity in isolated rat DRG neurons via 5-HT(4) receptors and cAMP-dependent mechanisms. Neuropharmacology 184, 108408 (2021).

76. R. Y. North, M. A. Odem, Y. Li, C. E. Tatsui, R. M. Cassidy, P. M. Dougherty, E. T. Walters, Electrophysiological Alterations Driving Pain-Associated Spontaneous Activity in Human Sensory Neuron Somata Parallel Alterations Described in Spontaneously Active Rodent Nociceptors. J Pain 23, 1343–1357 (2022).

77. M. A. Odem, A. G. Bavencoffe, R. M. Cassidy, E. R. Lopez, J. Tian, C. W. Dessauer, E. T. Walters, Isolated nociceptors reveal multiple specializations for generating irregular ongoing activity associated with ongoing pain. Pain 159, 2347–2362 (2018).

78. J. Tian, A. G. Bavencoffe, M. X. Zhu, E. T. Walters, Readiness of nociceptor cell bodies to generate spontaneous activity results from background activity of diverse ion channels and high input resistance. Pain 165, 893–907 (2024).

79. E. Velasco, J. L. Alvarez, V. M. Meseguer, J. Gallar, K. Talavera, Membrane potential instabilities in sensory neurons: mechanisms and pathophysiological relevance. Pain 163, 64–74 (2022).

80. S. Rose-John, Interleukin-6 Family Cytokines. Cold Spring Harb Perspect Biol 10, (2018).

81. C. Chipoy, M. Berreur, S. Couillaud, G. Pradal, F. Vallette, C. Colombeix, F. Redini, D. Heymann, F. Blanchard, Downregulation of osteoblast markers and induction of the glial fibrillary acidic protein by oncostatin M in osteosarcoma cells require PKCdelta and STAT3. J Bone Miner Res 19, 1850–1861 (2004).

82. S. E. Headland, H. S. Dengler, D. Xu, G. Teng, C. Everett, R. A. Ratsimandresy, D. Yan, J. Kang, K. Ganeshan, E. V. Nazarova, S. Gierke, C. J. Wedeles, R. Guidi, D. J. DePianto, K. B. Morshead, A. Huynh, J. Mills, S. Flanagan, S. Hambro, V. Nunez, J. E. Klementowicz, Y. Shi, J. Wang, J. Bevers, 3rd, V. Ramirez-Carrozzi, R. Pappu, A. Abbas, J. Vander Heiden, D. F. Choy, R. Yadav, Z. Modrusan, R. A. Panettieri, Jr., C. Koziol-White, W. F. Jester, Jr., B. J. Jenkins, Y. Cao, C. Clarke, C. Austin, D. Lafkas, M. Xu, P. J. Wolters, J. R. Arron, N. R. West, M. S. Wilson, Oncostatin M expression induced by bacterial triggers drives airway inflammatory and mucus secretion in severe asthma. Sci Transl Med 14, eabf8188 (2022).

83. C. Abbadie, S. Bhangoo, Y. De Koninck, M. Malcangio, S. Melik-Parsadaniantz, F. A. White, Chemokines and pain mechanisms. Brain research reviews 60, 125–134 (2009).

84. C. Sommer, M. Kress, Recent findings on how proinflammatory cytokines cause pain: peripheral mechanisms in inflammatory and neuropathic hyperalgesia. Neurosci Lett 361, 184–187 (2004).

85. M. S. Yousuf, S. I. Shiers, J. J. Sahn, T. J. Price, Pharmacological Manipulation of Translation as a Therapeutic Target for Chronic Pain. Pharmacological reviews 73, 59–88 (2021).

86. A. Yoshimura, T. Naka, M. Kubo, SOCS proteins, cytokine signalling and immune regulation. Nat Rev Immunol 7, 454–465 (2007).

87. K. Gomez, P. Duran, R. Tonello, H. N. Allen, L. Boinon, A. Calderon-Rivera, S. Loya-Lopez, T. S. Nelson, D. Ran, A. Moutal, N. W. Bunnett, R. Khanna, Neuropilin-1 is essential for vascular endothelial growth factor A-mediated increase of sensory neuron activity and development of pain-like behaviors. Pain 164, 2696–2710 (2023).

88. V. Jeevakumar, A. K. Al Sardar, F. Mohamed, C. M. Smithhart, T. Price, G. Dussor, IL-6 induced upregulation of T-type Ca(2+) currents and sensitization of DRG nociceptors is attenuated by MNK inhibition. J Neurophysiol 124, 274–283 (2020).

89. S. M. Mihail, A. Wangzhou, K. K. Kunjilwar, J. K. Moy, G. Dussor, E. T. Walters, T. J. Price, MNK-eIF4E signalling is a highly conserved mechanism for sensory neuron axonal plasticity: evidence from Aplysia californica. Philos Trans R Soc Lond B Biol Sci 374, 20190289 (2019).

90. D. Rodriguez, S. G. Kwatra, C. Dias-Barbosa, F. Zeng, Z. K. Jabbar Lopez, C. Piketty, J. Puelles, Patient Perspectives on Living With Severe Prurigo Nodularis. JAMA Dermatol 159, 1205–1212 (2023).

91. S. A. Bhuiyan, M. Xu, L. Yang, E. Semizoglou, P. Bhatia, K. I. Pantaleo, I. Tochitsky, A. Jain, B. Erdogan, S. Blair, V. Cat, J. M. Mwirigi, I. Sankaranarayanan, D. Tavares-Ferreira, U. Green, L. A. McIlvried, B. A. Copits, Z. Bertels, J. S. Del Rosario, A. J. Widman, R. A. Slivicki, J. Yi, C. J. Woolf, J. K. Lennerz, J. L. Whited, T. J. Price, R. W. Gereau, W. Renthal, Harmonized cross-species cell atlases of trigeminal and dorsal root ganglia. bioRxiv, (2023).

92. S. Shiers, R. M. Klein, T. J. Price, Quantitative differences in neuronal subpopulations between mouse and human dorsal root ganglia demonstrated with RNAscope in situ hybridization. Pain 161, 2410–2424 (2020).

93. S. Shiers, I. Sankaranarayanan, V. Jeevakumar, A. Cervantes, J. C. Reese, T. J. Price, Convergence of peptidergic and non-peptidergic protein markers in the human dorsal root ganglion and spinal dorsal horn. bioRxiv, 2020.2010.2014.339382 (2020).

94. R. V. Haberberger, C. Barry, N. Dominguez, D. Matusica, Human Dorsal Root Ganglia. Front Cell Neurosci 13, 271 (2019).

95. D. C. Molliver, D. E. Wright, M. L. Leitner, A. S. Parsadanian, K. Doster, D. Wen, Q. Yan, W. D. Snider, IB4-binding DRG neurons switch from NGF to GDNF dependence in early postnatal life. Neuron 19, 849–861 (1997).

96. S. E. Jordt, D. Julius, Molecular basis for species-specific sensitivity to "hot" chili peppers. Cell 108, 421–430 (2002).

97. W. J. Laursen, E. R. Schneider, D. K. Merriman, S. N. Bagriantsev, E. O. Gracheva, Low-cost functional plasticity of TRPV1 supports heat tolerance in squirrels and camels. Proc Natl Acad Sci U S A 113, 11342–11347 (2016).

98. W. Jiang, J. D. Glaeser, G. Kaneda, J. Sheyn, J. T. Wechsler, S. Stephan, K. Salehi, J. L. Chan, W. Tawackoli, P. Avalos, C. Johnson, C. Castaneda, L. E. A. Kanim, T. Tanasansomboon, J. E. Burda, O. Shelest, H. Yameen, T. G. Perry, M. Kropf, J. M. Cuellar, D. Seliktar, H. W. Bae, L. S. Stone, D. Sheyn, Intervertebral disc human nucleus pulposus cells associated with back pain trigger neurite outgrowth in vitro and pain behaviors in rats. Sci Transl Med 15, eadg7020 (2023).

99. W. Renthal, A. Chamessian, M. Curatolo, S. Davidson, M. Burton, S. Dib-Hajj, P. M. Dougherty, A. D. Ebert, R. W. T. Gereau, A. Ghetti, M. S. Gold, G. Hoben, D. M. Menichella, P. Mercier, W. Z. Ray, D. Salvemini, R. P. Seal, S. Waxman, C. J. Woolf, C. L. Stucky, T. J. Price, Human cells and networks of pain: Transforming pain target identification and therapeutic development. Neuron 109, 1426–1429 (2021).

100. I. Sankaranarayanan, D. Tavares-Ferreira, J. M. Mwirigi, G. L. Mejia, M. D. Burton, T. J. Price, Inducible co-stimulatory molecule (ICOS) alleviates paclitaxel-induced neuropathic pain via an IL-10-mediated mechanism in female mice. J Neuroinflammation 20, 32 (2023).

