## Supplementary Figures for "Expansion of OSMR expression and signaling in the human dorsal root ganglion links OSM to neuropathic pain"

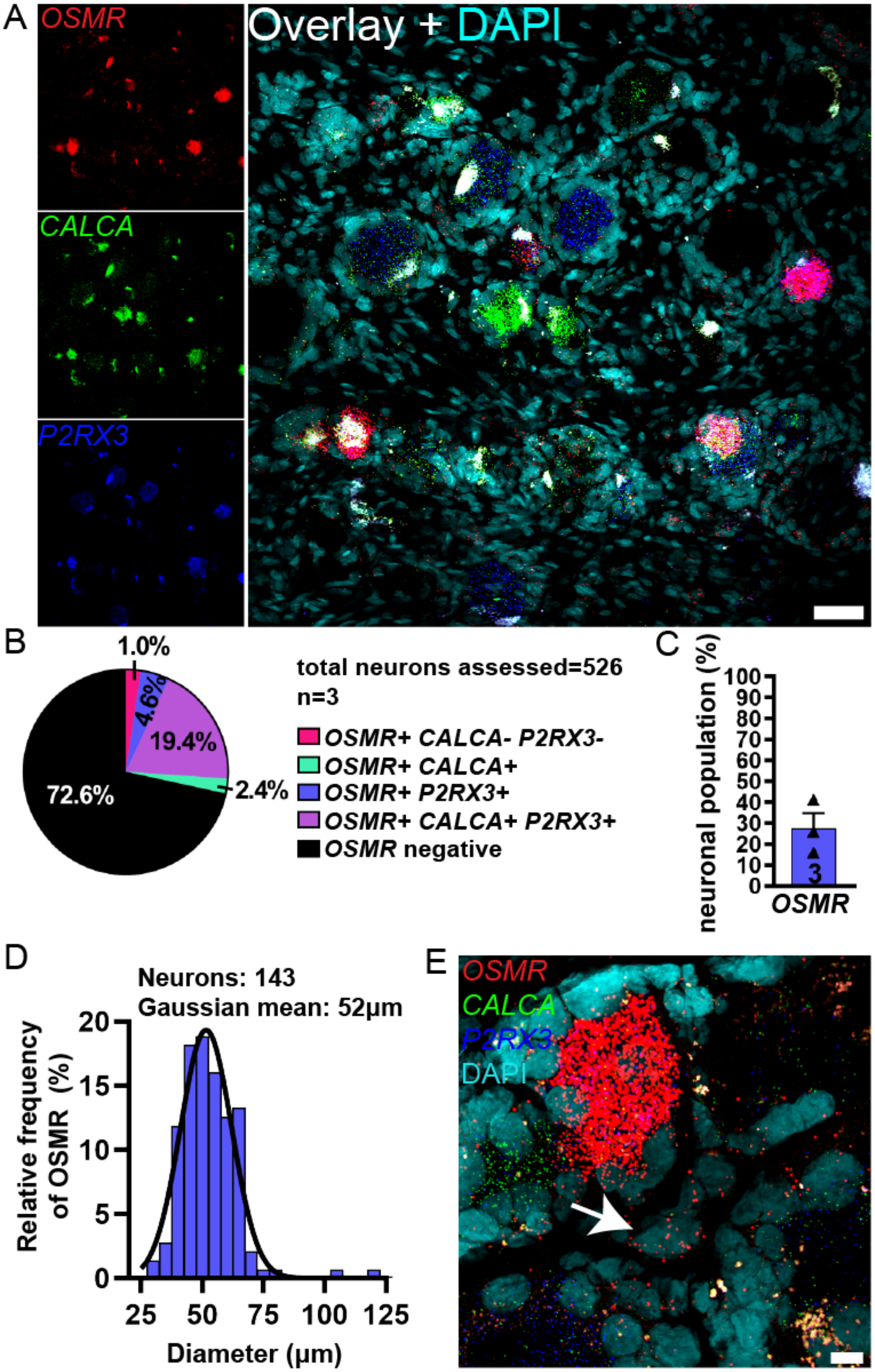
A
OSMR
CALCA
P2RX3
Overlay + DAPI
B
1.0%
4.6%
19.4%
2.4%
72.6%
total neurons assessed=526
n=3
OSMR+ CALCA- P2RX3-
OSMR+ CALCA+
OSMR+ P2RX3+
OSMR+ CALCA+ P2RX3+
OSMR negative
C
neuronal population (%)
100
90
80
70
60
50
40
30
20
10
0
3
OSMR
D
Neurons: 143
Gaussian mean: 52μm
Relative frequency of OSMR (%)
20
15
10
5
0
25
50
75
100
125
Diameter (μm)
E
OSMR
CALCA
P2RX3
DAPI

**Fig. S1. General distribution of *OSMR* mRNA in human dorsal root ganglia. A**) Representative 20x images of human dorsal root ganglion (hDRG) labeled with RNAscope *in situ* hybridization for *CALCA* (green), *P2RX3* (blue), and *OSMR* (red) mRNA and costained with DAPI (cyan). Scale bar, 50 µm. Lipofuscin (globular structures) that autofluoresced in all 3 channels and appear white in the overlay image were not analyzed as this is background signal that is present in all human nervous tissue. **B**) Pie chart displaying the distribution of *OSMR* neuronal subpopulations in the human DRG. Pie chart is an average of all 3 donors. **C**) *OSMR* was expressed in 27.4% of all sensory neurons in human DRG. **D**) Histogram with a fitted Gaussian distribution displaying the diameter measurements of all *OSMR*-positive neurons between all 3 donors. **E**) A representative 100x overlay image demonstrates that *OSMR* is also highly expressed in non-neuronal cells, as the white arrow exemplifies. These cells created a ring-like shape around the neurons, a pattern indicative of satellite glial cells. Scale bar, 10 µm.

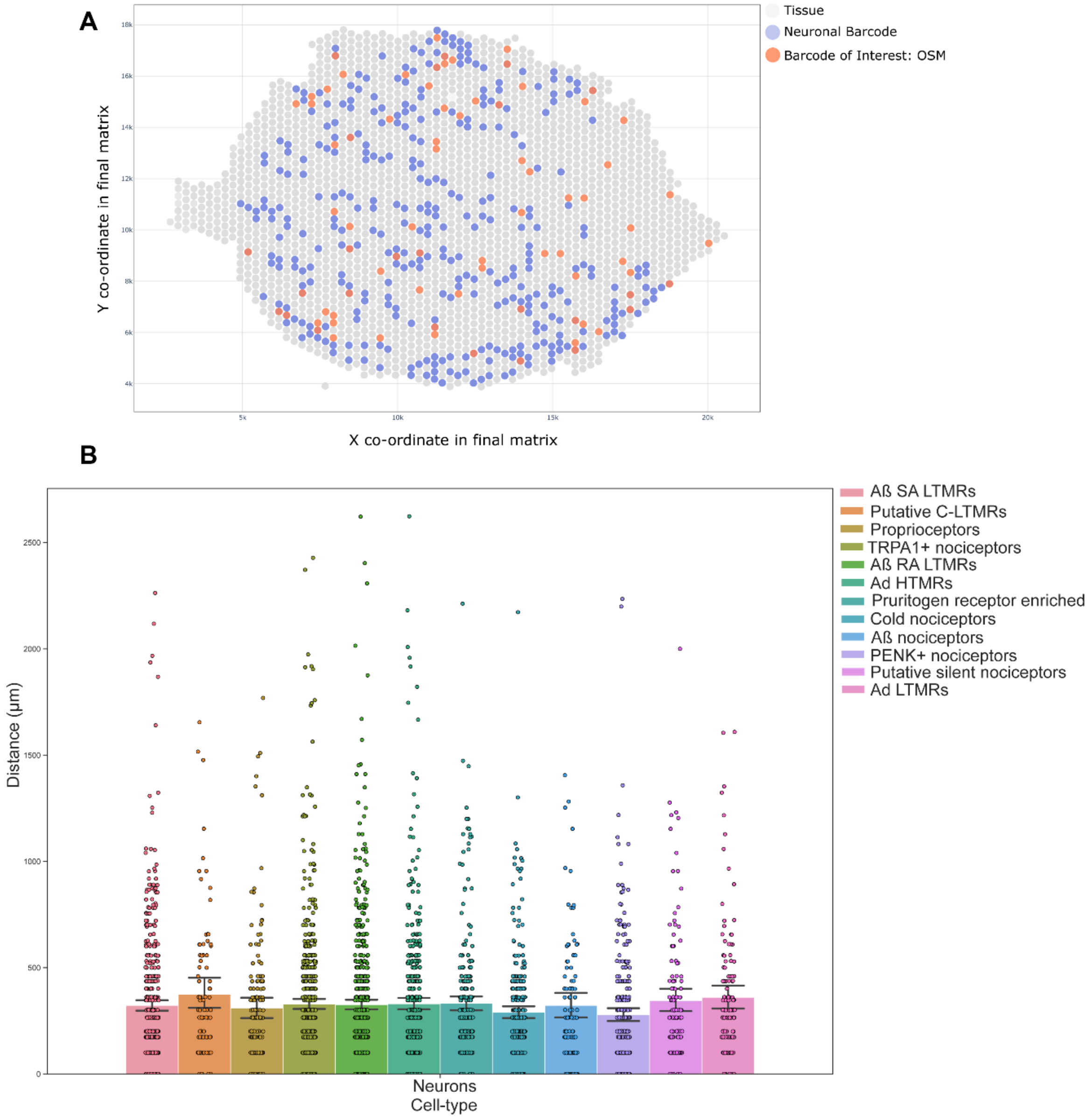


**Fig. S2. Spatial transcriptomics of human DRG shows that OSM+ barcodes are in close proximity to neurons in the human DRG. A**) Spatial distribution of OSM positive barcodes and their proximity to neuronal barcodes in human dorsal root ganglia (hDRG). **B**) The minimum distance of an OSM+ barcode to each neuronal subtypes (in µm).

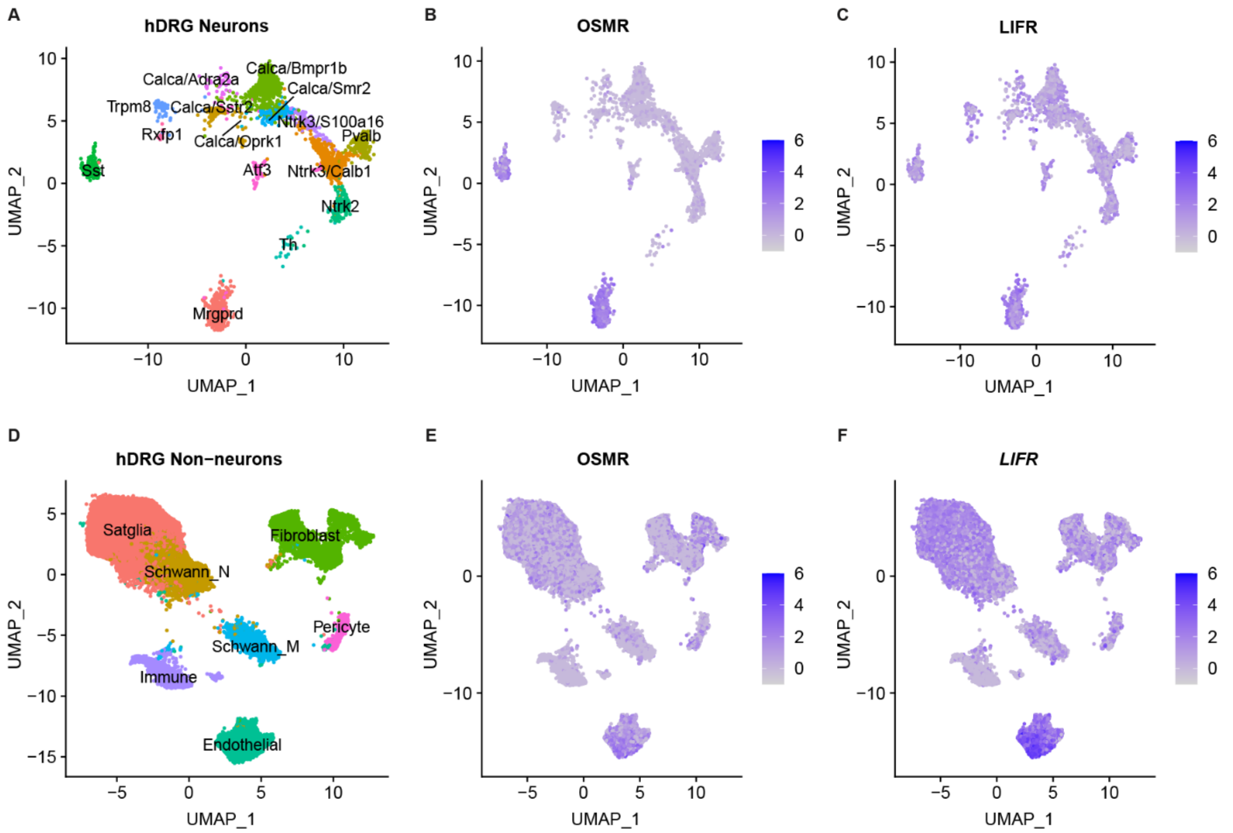


**Fig. S3. *LIFR* is more abundantly expressed across human DRG neurons and non-neuronal cell types than *OSMR*.** Transcriptomic sequencing data are from the harmonized DRG atlas by Bhuiyan et al., 2024, and only show hDRG populations. **A**) UMAP plot highlighting distinct neuronal subtypes in hDRG neurons based on their respective markers, labeled using mouse nomenclature. UMAP plots illustrating the expression of *OSMR* **B**) and *LIFR* **C**) across hDRG neurons, demonstrating differential expression patterns. **D**) UMAP plot showing clustering of hDRG non-neuronal cell types. UMAP plots depict the expression of *OSMR* **E**) and *LIFR* **F**) across non-neuronal cell populations in the hDRG. *LIFR* shows more widespread and higher expression levels than *OSMR* across both neuronal and non-neuronal cell types, suggesting it plays a broader role in hDRG function.

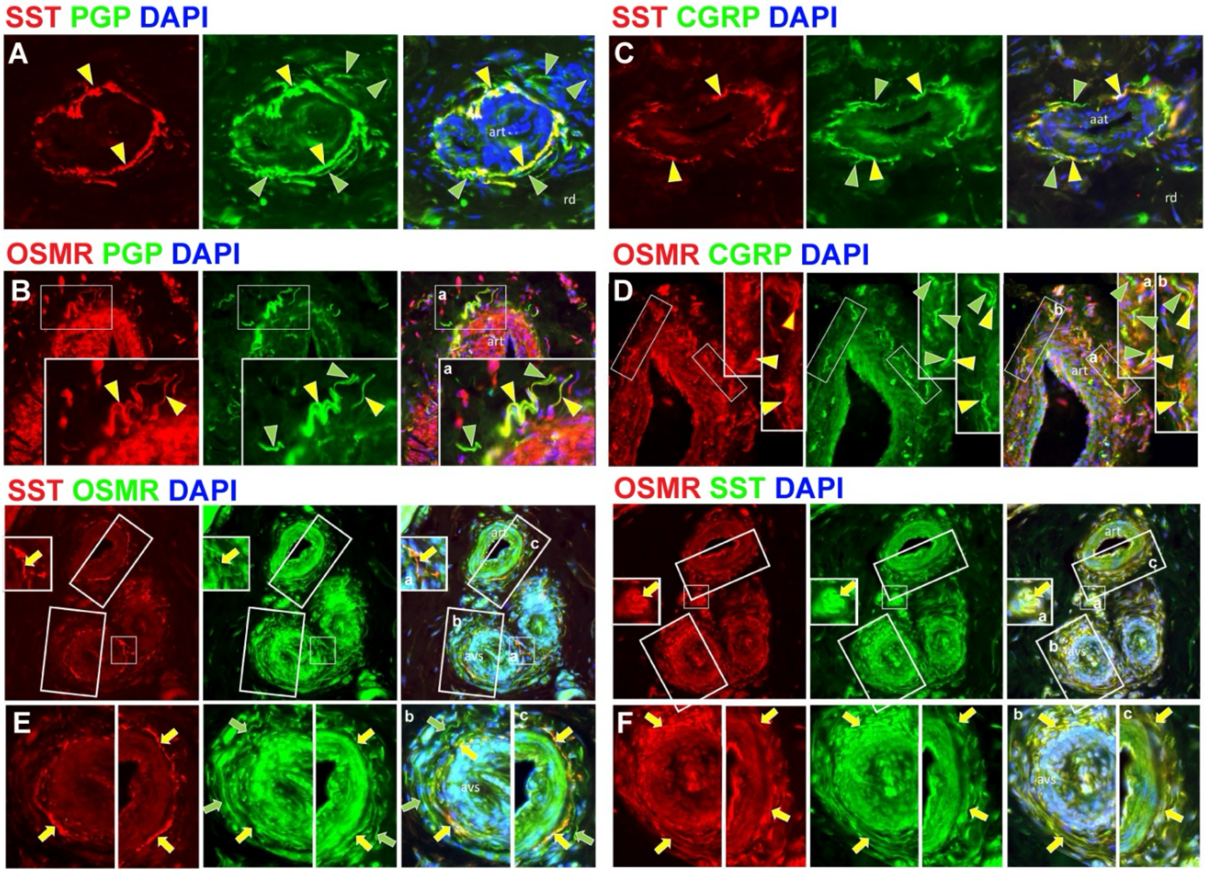


**Fig. S4 Immunofluorescence labeling of human palmar skin biopsies reveals stratified subtypes of OSMR only, SST only, as well as OSMR and SST coexpressing innervation of arterioles (art) and arterio-venule shunts (AVS) located in the deep reticular dermis (rd).** All images, lettering, and symbols are as described in and are at the same magnification as Figure 2. The wall of the arterioles and AVS consists of the inner endothelial cell lining (ec), thick smooth muscle tunica medularis (tm), and outer connective tissue tunica adventitia (ta). All of the innervation terminates in the tunica adventitia. **A, C**) SST and OMSR labeling (yellow arrowheads) is respectively on subsets among the total innervation that labels for PGP9.5. Noradrenergic sympathetic innervation is known to be among that only labeled for PGP9.5 (green arrowheads**. B, D**) All the SST and OMSR are co-expressed among CGRP sensory innervation (yellow arrowheads), which is known to be the largest contingent of the sensory innervation. Note that SST is expressed virtually only on innervation, whereas OSMR is also expressed on endothelial cells, smooth muscle cells, and other scattered cells in the dermis (small white arrows). Some CGRP innervation lacks SST and OSMR (green arrowheads). **E, F**) Reverse order label combinations taken together with **A-D**) indicate that OMSR is expressed without SST among the innervation to outer tunica adventitia, whereas innervation concentrated near the tunica muscularis extensively co expresses SST and OMSR.

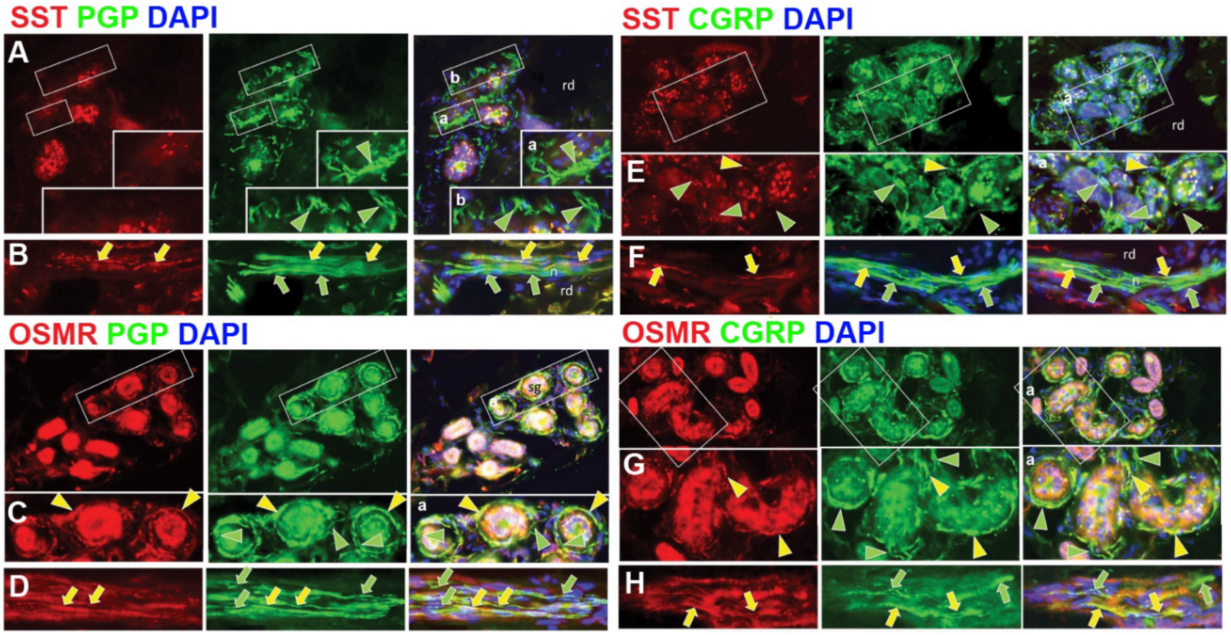


**Fig. S5 Immunofluorescence labeling of human palmar skin biopsies reveals only OSMR expression on sweat gland (SG) innervation and SST and/or OSMR expression among axons in small nerves located in the deep reticular dermis (rd).** All images, lettering, and symbols are as described in and are at the same magnification as Figure 2. **A,B,E,F**) SST labeling (yellow arrowhead) is rarely seen among all the innervation to sweat glands (green arrowheads). SST labeling (yellow arrows) was co-expressed on many but not all (green arrows) axons that are CGRP+. **C,D,G,H**) OSMR (yellow arrowheads) is expressed among many, but not all (green arrowheads) of the CGRP innervation to sweat glands. OSMR labeling (yellow arrows) was co-expressed on many but not all (green arrows) that are CGRP+.

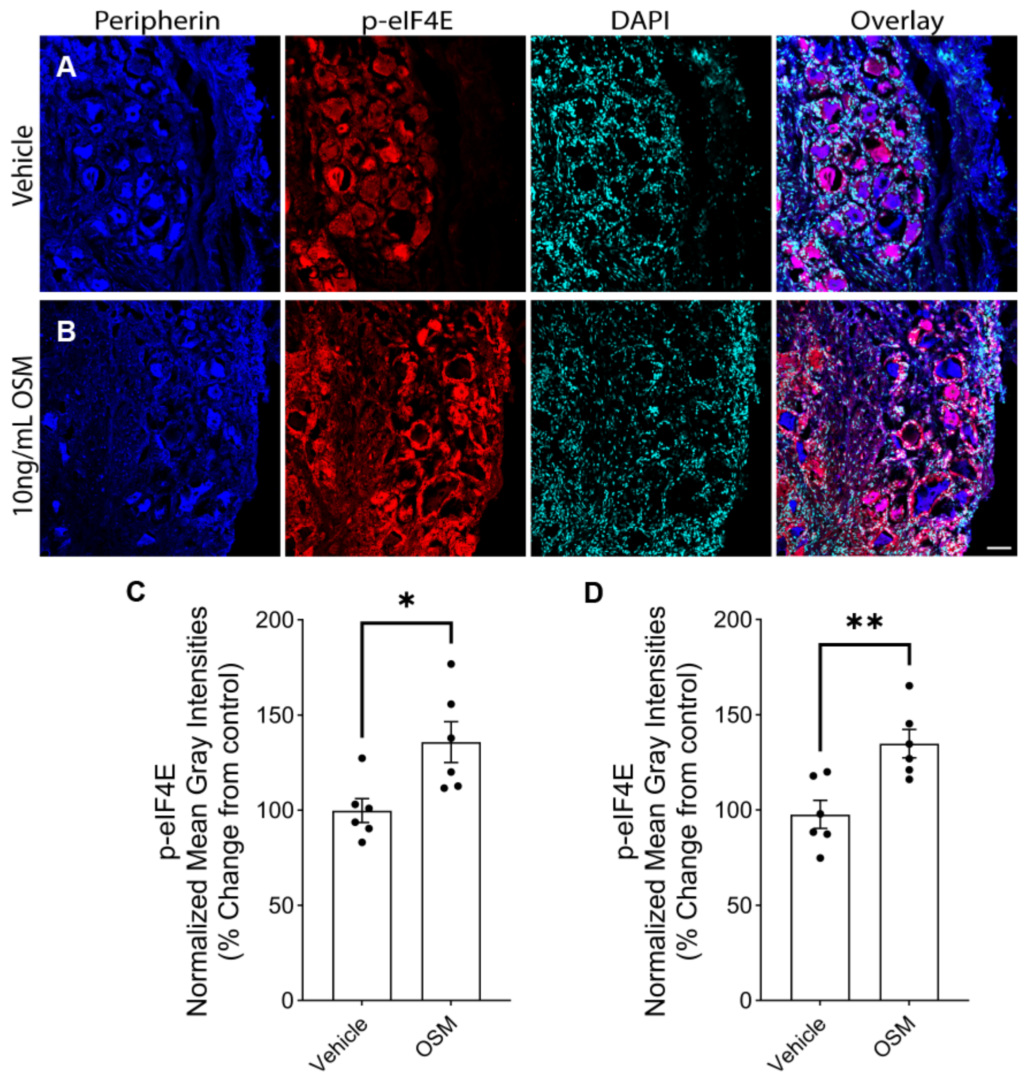


**Fig. S6. OSM induces MNK- eIF4E signaling in hDRG explants. A** and **B**) peripherin-labeled neurons show significant increases in the phosphorylation of eIF4E (p-eIF4E, red) signal intensity in **C**) neurons and **D**) surrounding glial-like cells of OSM-treated hDRG explants. DAPI (cyan) labels all nuclei. 20X Magnification, Scale bar, 50µm. N=6 technical replicates from 2 organ donors. Unpaired t-test *p<0.05, **p<0.01.

**A**

**Primer sequences for OSMR and LIFR CRISPR KO validation**

| Gene | Sequence | | $T_m$ (ºC) | $T_a$ (ºC) | bp |
|---|---|---|---|---|---|
| OSMR | Forward | 5’- TCCCAACCGGTTAGATTGTC -3’ | 51.406 | 58 | 682 |
| | Reverse | 5’- GTGCATCCCATCCTACCTAC -3’ | 51.349 | 58 | |
| LIFR | Forward | 5’- CCAGGGAAGCTTGAGTTTGA -3’ | 51.879 | 58 | 614 |
| | Reverse | 5’- TCTTGATTGTGCTGGTGGTT -3’ | 51.761 | 58 | |

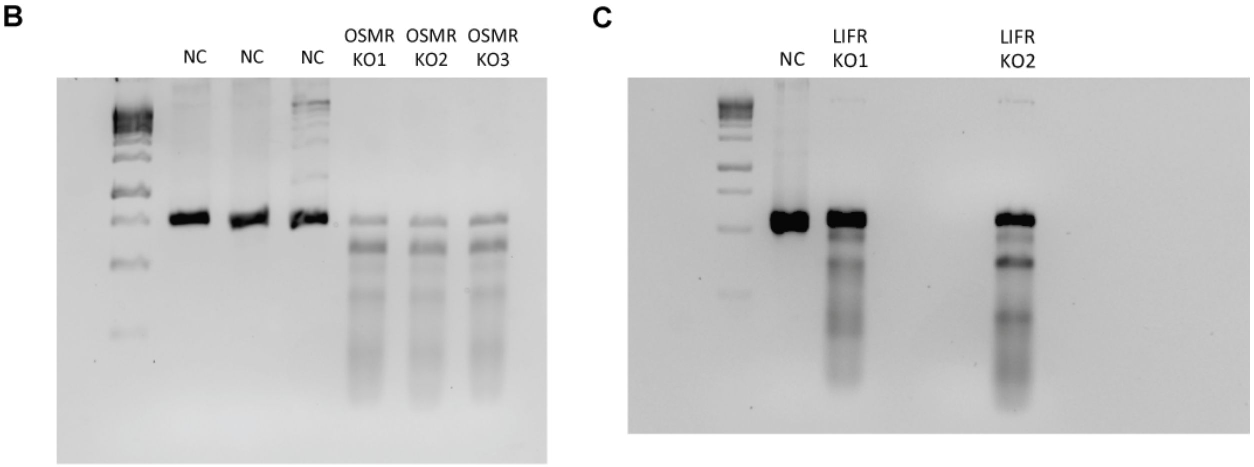


**Fig. S7. Validation of CRISPR knockout. A**) Forward and reverse primers used for the validation of OSMR and LIFR knockout. **B, C**) Representative PCR results confirming genomic editing at the OSMR and LIFR loci, respectively. Negative control (NC) and knockout (KO) bands are indicated, with expected band sizes of ~682 bp for OSMR and ~614 bp for LIFR.

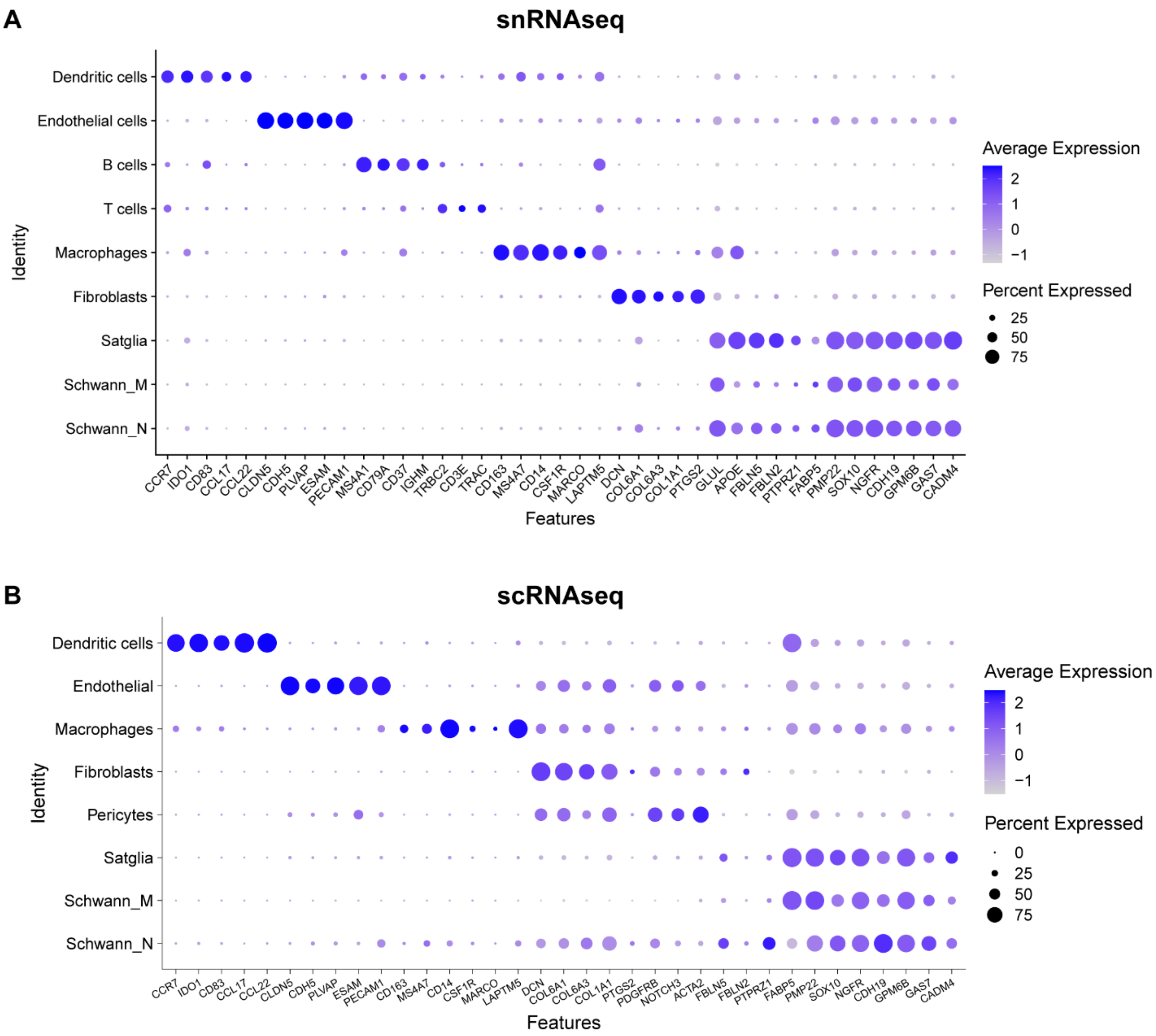


**Fig. S8. Dot plots of non-neuronal population markers in the A) snRNAseq and B) scRNAseq datasets from OSM- and Vehicle- treated hDRG cultured cells.** Dot size indicates the fraction of expressing cells, and color intensity reflects the average expression level.

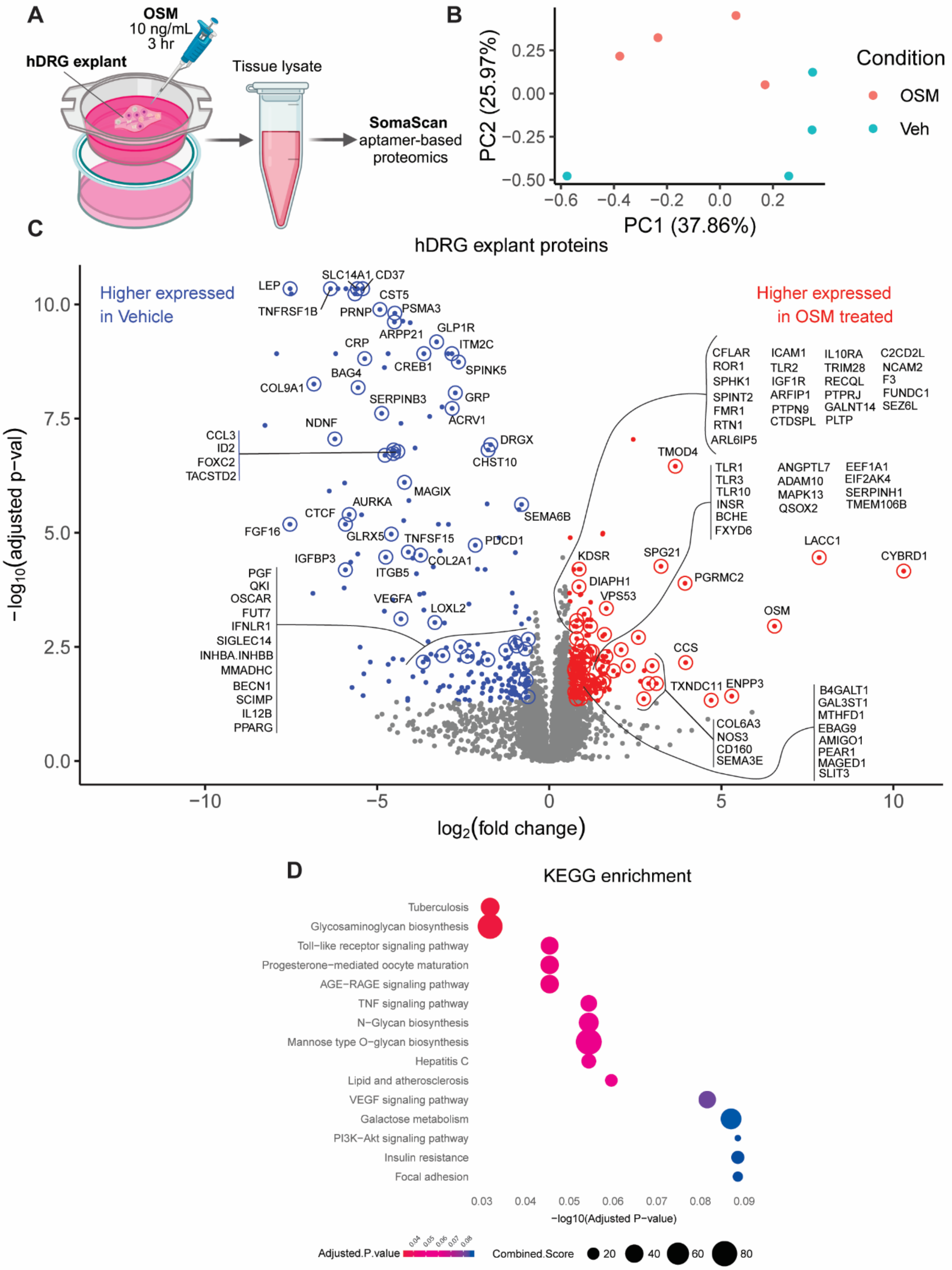

A
OSM
10 ng/mL
3 hr
hDRG explant
Tissue lysate
SomaScan
aptamer-based
proteomics
B
PC2 (25.97%)
PC1 (37.86%)
Condition
OSM
Veh
C
hDRG explant proteins
Higher expressed
in Vehicle
Higher expressed
in OSM treated
−log10(adjusted p−val)
log2(fold change)
D
KEGG enrichment
Tuberculosis
Glycosaminoglycan biosynthesis
Toll−like receptor signaling pathway
Progesterone−mediated oocyte maturation
AGE−RAGE signaling pathway
TNF signaling pathway
N−Glycan biosynthesis
Mannose type O−glycan biosynthesis
Hepatitis C
Lipid and atherosclerosis
VEGF signaling pathway
Galactose metabolism
PI3K−Akt signaling pathway
Insulin resistance
Focal adhesion
−log10(Adjusted P−value)
Adjusted.P.value
Combined.Score

**Fig. S9. OSM induces distinct proteomic changes in hDRG explants. A)** Experimental design: hDRG explants were treated with 10ng/mL OSM or Vehicle for 3 hours. Following treatment, we extracted tissue lysates and performed SomaScan aptamer-based proteomic profiling to measure changes in protein expression. **B)** PCA plot illustrates the shift in overall protein expression driven by OSM. **C)** Volcano plot of differentially expressed proteins (labeled by gene names). Differentially upregulated proteins ($Log_2FC \geq 0.585$, adjusted p-value $< 0.05$) are shown in red, and downregulated proteins ($Log_2FC \leq -0.585$) are shown in blue. **D)** KEGG pathway enrichment analysis. Proteins that change significantly in response to OSM treatment were mapped to KEGG pathways, highlighting major biological processes and signaling pathways modulated by OSM in hDRG explants.

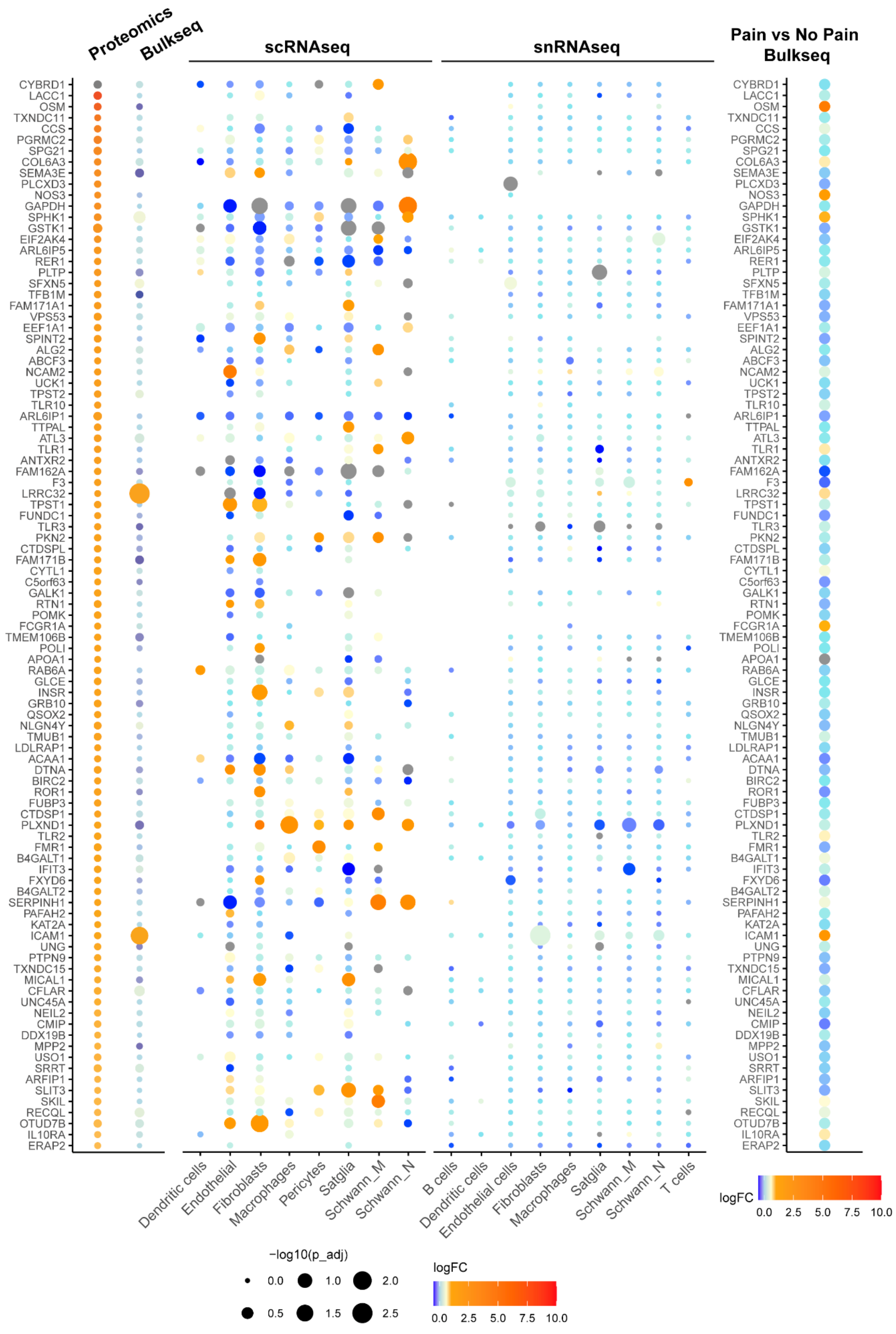

Proteomics
Bulkseq
scRNAseq
snRNAseq
Pain vs No Pain Bulkseq
CYBRD1
LACC1
OSM
TXNDC11
CCS
PGRMC2
SPG21
COL6A3
SEMA3E
PLCXD3
NOS3
GAPDH
SPHK1
GSTK1
EIF2AK4
ARL6IP5
RER1
PLTP
SFXN5
TFB1M
FAM171A1
VPS53
EEF1A1
SPINT2
ALG2
ABCF3
NCAM2
UCK1
TPST2
TLR10
ARL6IP1
TTPAL
ATL3
TLR1
ANTXR2
FAM162A
F3
LRRC32
TPST1
FUNDC1
TLR3
PKN2
CTDSPL
FAM171B
CYTL1
C5orf63
GALK1
RTN1
POMK
FCGR1A
TMEM106B
POLI
APOA1
RAB6A
GLCE
INSR
GRB10
QSOX2
NLGN4Y
TMUB1
LDLRAP1
ACAA1
DTNA
BIRC2
ROR1
FUBP3
CTDSP1
PLXND1
TLR2
FMR1
B4GALT1
IFIT3
FXYD6
B4GALT2
SERPINH1
PAFAH2
KAT2A
ICAM1
UNG
PTPN9
TXNDC15
MICAL1
CFLAR
UNC45A
NEIL2
CMIP
DDX19B
MPP2
USO1
SRRT
ARFIP1
SLIT3
SKIL
RECQL
OTUD7B
IL10RA
ERAP2
Dendritic cells
Endothelial
Fibroblasts
Macrophages
Pericytes
Satglia
Schwann_M
Schwann_N
B cells
Dendritic cells
Endothelial cells
Fibroblasts
Macrophages
Satglia
Schwann_M
Schwann_N
T cells
logFC
0.0
2.5
5.0
7.5
10.0
−log10(p_adj)
0.0
0.5
1.0
1.5
2.0
2.5

**Fig. S10. Comparison of the top 100 significantly upregulated proteins with multiple transcriptomic datasets reveals shared and distinct expression changes.** These proteins, identified by proteomic analysis of OSM-treated hDRG explants, were cross-referenced with four RNAseq datasets: (1) bulk RNAseq of OSM-treated hDRG cultures, (2) scRNAseq), (3) snRNAseq, and (4) bulk RNAseq from neuropathic pain patients (pain vs. no-pain DRGs). The integrated analysis highlights cell type–specific gene and protein expression changes, revealing both overlapping and distinct pathways influenced by OSM treatment and neuropathic pain.
